# Long-term retention of antigens in germinal centres is controlled by the spatial organisation of the follicular dendritic cell network

**DOI:** 10.1101/2022.09.06.506650

**Authors:** Ana Martinez-Riano, Shenshen Wang, Stefan Boeing, Sophie Minoughan, Antonio Casal, Katelyn M Spillane, Burkhard Ludewig, Pavel Tolar

## Abstract

Germinal centers (GCs) require sustained availability of antigens to promote antibody affinity maturation against pathogens and vaccines. A key source of antigens for GC B cells are immune complexes (ICs) displayed on follicular dendritic cells (FDCs). Here we show that FDC spatial organization regulates antigen dynamics in the GC. We show the existence of two light zone (LZ) FDC populations, which differ in the duration of antigen retention. While the entire light zone (LZ) FDC network captures ICs initially, only the central cells of the network function as a long-term antigen reservoir, where different antigens arriving from subsequent immunizations co-localize. Mechanistically, central FDCs constitutively express subtly higher CR2 membrane densities than peripheral FDCs, which strongly increases the IC retention half-life. Even though repeated immunizations gradually saturate central FDCs, B cell responses remain efficient because new antigens partially displace old ones. These results reveal the principles shaping antigen display on FDCs during the GC reaction.

## INTRODUCTION

The generation of high-affinity antibodies that recognise and neutralise pathogens is a key hallmark of the humoral response. The response is initiated by naïve B cells located in the follicles of the secondary lymphoid organs, which are constantly exposed to new antigens arriving from the lymph or blood. B cells activated by their cognate antigen generate an initial burst of antibody-secreting and memory cells and also seed germinal centers (GCs), where B cell receptor (BCR) diversification and selection for antigen affinity takes place(Victora and Nussenzweig, 2022). In GCs, B cells with higher affinity BCRs outcompete B cells with lower affinity BCRs thanks to the survival and proliferative advantage instigated by higher binding to antigens, higher antigen presentation on MHC-II and higher T cell help(Gitlin et al., 2014; Victora et al., 2010). This selection takes several weeks and results in an increase in the affinity of the antibodies produced by GC-generated plasma cells (PCs). Thus, a constant supply of antigens to GC B cells is crucial to enable affinity maturation critical for antibody-mediated protection.

The entry of antigens into B cell follicles and their retention are controlled by the specific cellular organisation of secondary lymphoid organs(Batista and Harwood, 2009; Mueller and Germain, 2009). The cellular subset most closely associated with antigen retention in GCs are follicular dendritic cells (FDCs)(Alexandre et al., 2022; Pikor et al., 2020; Rodda et al., 2018). FDCs are stromal cells that develop from perivascular or subcapsular precursors of the spleen and lymph node, respectively(Jarjour et al., 2014; Krautler et al., 2012) in response to LTA1B2 and TNF produced by B cells(Hir et al., 1996; Pasparakis et al., 1996; Tumanov et al., 2004; Wang et al., 2001). Mature FDCs form tight networks via long intermingled dendrites and control follicle homeostasis thanks to the expression of cytokines and factors that regulate B cell survival, localisation and elimination upon apoptosis (e. g. CXCL13(Wang et al., 2011), GGT5(Lu et al., 2019), TNFSF13B (BAFF)(Suzuki et al., 2010) and MFGE8(Kranich et al., 2008)). Recently, a subset of FDCs was also shown to produce CXCL12 and orchestrate the dark zone (DZ) of the GC(Pikor et al., 2020; Rodda et al., 2015).

FDCs bind antigens in the form of particles or immune complexes (ICs) coated by complement. Within the first day after immunisation, FDCs acquire the complement-coated antigens from non-cognate B cells, which shuttle the antigen from the subcapsular sinus of the LN or from the marginal zone of the spleen(Ferguson et al., 2004; Phan et al., 2007). Subsequently, FDCs are the only cells known to keep and display the antigens in their native conformation for long periods of time, thanks to the internalisation of the antigens into non-degradative endosomes and their recycling to the cell surface(Heesters et al., 2013).

On the molecular level, FDCs bind the antigens coated by the C3d complement fragment using two complement receptors CR1 (CD35) and CR2 (CD21). In the mouse, both receptors are encoded by a single *Cr2* gene and we, therefore, refer to them both as CR2 here. In addition, FDCs also express an array of Fc-receptors (FCGR2B, FCER2A and FCMAR)(Allen and Cyster, 2008), whose role is less clear, but may be important in secondary or autoimmune responses(Gonzalez et al., 2010; Poel et al., 2019; Shikh et al., 2006; Tokatlian et al., 2019). Thus, FDC-retained antigens are particularly important for the sustained production of antibodies and for secondary responses, possibly by prolonging the production of antigenspecific PCs and memory B cells in GCs. However, in contrast to the initial capture of antigens, the dynamics of FDC antigen retention over the GC response and secondary immunizations remain poorly understood. Similarly, it is unclear whether the precise topological organization of FDCs in the follicle underpins their functional relevance(Allen and Cyster, 2008). To better understand the long-term antigen dynamics on FDCs after primary and secondary immunizations, we imaged clarified LNs from mice immunised with fluorescent antigen-ICs over time. We observed striking differences in antigen retention functionality of the FDCs based on the localization of the cells. While the entire FDC network contributes to the initial antigen capture and display to B cells, only the central cells of the network participate in the long-term antigen retention in the GC and serve as antigen reservoirs. Importantly, the topological organization of the network also influence subsequent antigens, which follow the same kinetics and are retained centrally over time. GC-derived signals contribute to the FDC network expansion and maturation but do not regulate the centralization of the antigens. We observed that the differences in antigen display over time are the consequence of different levels of CR2 expression by the peripheral and central FDCs of the network. The higher CR2 levels in the central FDCs prevent fast dissociation of the antigen-ICs and increase their interaction half-life. Single-cell transcriptomics corroborated the FDC heterogeneity and suggested that peripheral and central FDCs are functionally distinct populations among light zone (LZ) FDCs. Finally, we observed that repeated immunizations partially saturate the central FDCs and partially displace old antigens by new ones. Nevertheless, the efficiency of antigen presentation to B cells by FDCs is sufficient for antibody production. Understanding the basis of antigen dynamics on the FDC network will guide the generation of more efficient vaccines through the improvement of antigen retention in the GC.

## RESULTS

### Long-term antigen retention in B cell follicles is performed by a subpopulation of FDCs residing in the follicle center

To study antigen availability to B cells during the GC response, we immunised mice with fluorescent antigen immune complex (IC) and analyzed draining lymph nodes (LN) at different time-points post-injection (Figure 1A). Immunocomplexed antigens elicit complement deposition, traffic efficiently into the B cell follicles and bind to FDCs(Phan et al., 2007). To obtain a full view of antigen localization within the entire LN, we clarified the LNs, stained them with anti-CR2 antibodies to label FDCs and imaged them by 3D confocal microscopy. We observed that after 24 hours postimmunisation, most of the antigen was loaded onto the entire FDC network of each B cell follicle. However, on days 7 and 14 after immunization, the localisation of the antigen became restricted to the centers of each FDC network (Figure 1B and Suppl. Figure 1A). To quantify the distribution of the antigen within the FDC network over time, we generated an image analysis workflow which detected the FDC network of each follicle in 3D and divided them into concentric shells (Suppl. Figure 1B). The antigen quantity per FDC in each shell was quantified as the ratio of antigen to anti-CR2 fluorescence. The quantification showed that on day 1 postimmunization, antigen was distributed equally across the follicle (Figure 1C). However, 7 days after immunization, the antigen was preferentially located in the center of the FDC network, and a similar distribution was detected on day 14 (Figure 1C). The apparent centralization of the antigen was independent of the presence of adjuvant during immunization, since it was similar when the antigen was injected in alum or in PBS (Figure 1C, D, Suppl. Figure 1C, D).

**Figure 1.**
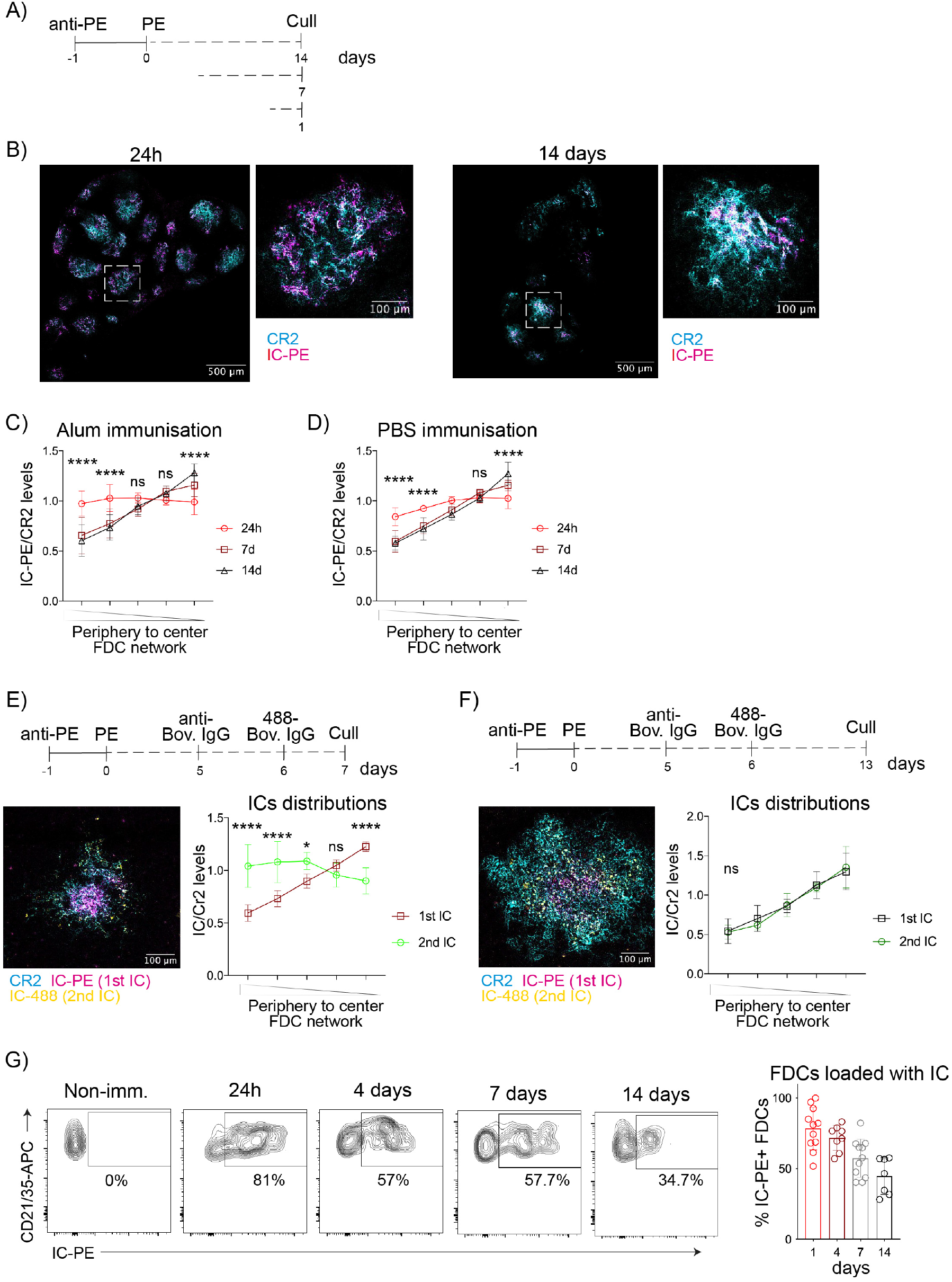
Long-term retention of antigens in B cell follicles takes place in the center of the FDC network. A) Immunization workflow to analyse IC-PE deposition on FDCs after 24 hours, 7 and 14 days. IC-PE immunization is performed by injecting PE-specific antibody (anti-PE) i. p. followed by PE s.c. the day after. B) Confocal images of clarified draining LNs 24 hours (left) and 14 days (right) after immunization with IC-PE (magenta). FDC networks are stained by anti-CD21/35 antibody binding to CR2 on FDCs (cyan). White squares indicate the region magnified. C) Quantification of antigen distribution on the FDC networks of mice immunized with IC-PE for 24 hours, 7 and 14 days with antigen embedded in Alum. Quantification workflow is described in Suppl. Figure 1A (Symbols show means ± SD from 6-9 LN, aggregated from 6-9 mice in 3 experiments). D) Antigen distribution on the FDC network of mice immunized as described in (C) with antigen diluted in PBS (n = 6-7 LN, 2 experiments). E) Distribution of ICs on the FDC network 1 day after a repeated immunization. LNs were analyzed 7 days after the first IC injection (IC-PE) and 24 hours after the second one (IC-488). Left, a representative confocal image of an FDC network of one follicle (anti-CD21/35 in cyan; IC-PE in magenta; IC-488 in yellow). Right, graph showing the quantification of the distribution of both ICs in the FDC network (n = 6 LN from 2 experiments). F) Distribution of ICs on the FDC network 7 days after the repeated immunization (IC4-88, 2^nd^ IC) and 14 days (IC-PE, 1^st^ IC) after the first immunization as described for (E) (n = 5-7 LN from 5-7 mice, 2 experiments). G) Flow cytometry plots and quantification of the percentage of IC-PE^+^ FDCs at different time-points after IC-PE immunization (n = 7-11 mice). Quantitative data shows means ± SD. Two-way’s ANOVA with multiple comparisons, ns, P > 0.05; *, P < 0.05; **, P < 0.01; ** *, P < 0.001; ** **, P < 0.0001

To understand if subsequent immunizations generate a similar pattern of antigen distribution, we immunized mice with a first fluorescent antigen-IC (IC-PE) and 6 days later with a second, different antigen-IC (IC-488; Figure 1E and F). We imaged clarified draining LNs at 24 hours and 7 days after the second immunization and analyzed the localization of both antigens. The imaging showed that 24 hours after the second immunisation, the second antigen-IC (IC-488) was present over the entire FDC network, while the first antigen was already in the center. 7 days later, IC-488 distribution changed and overlapped with the first antigen, IC-PE, on the central FDCs of each follicle. Thus, while the entire FDC network captures incoming ICs initially, the retention of the antigens takes place exclusively on the central FDCs of the network, independently of the presence of previous antigens.

Antigen retention on the central FDCs could be observed up to 56 days post-immunisation (Suppl. Figure 1E). On day 21, the location of the antigen overlapped with GCs (Suppl. Figure 1F). In addition, selective retention of antigens in the center of the FDC network was also observed 7 days after immunization of mice with HIV-gp120 nanoparticles (Suppl. Figure 1G), a vaccine immunogen shown to trigger complement activation independently of IC formation(Tokatlian et al., 2019). Furthermore, flow cytometry analysis of FDCs extracted from the LNs of mice immunised with fluorescent antigen-IC corroborated the imaging, showing that the percentage of FDCs loaded with antigen-IC after immunisation decreased over time (Suppl. Figure 1H, Figure 1G). These data suggested the presence of two subpopulations of FDCs, peripheral and central, that are both capable of capturing antigens transiently but differ in their retention of the antigens beyond the first week post-immunization.

### The expansion of FDCs induced by GC-derived signals is not responsible for the changes in antigen distribution

Changes in antigen localization may be driven by reorganization of the FDC network after immunization. FDC development and maintenance continuously depend on signalling induced by LTA1B2 and TNF, which are produced by B cells and whose production increases after B cell activation(Ansel et al., 2000; Endres et al., 1999; Fu et al., 1998; Gonzalez et al., 1998). We observed that in mice immunised with IC, the number of FDCs in the draining LNs increased compared to non-immunised mice (Figure 2A). To understand if changes in the FDC network cellularity could be responsible for the dynamics of antigen localization and retention, we analysed mice in which antigen-dependent B cell activation is impaired.

**Figure 2.**
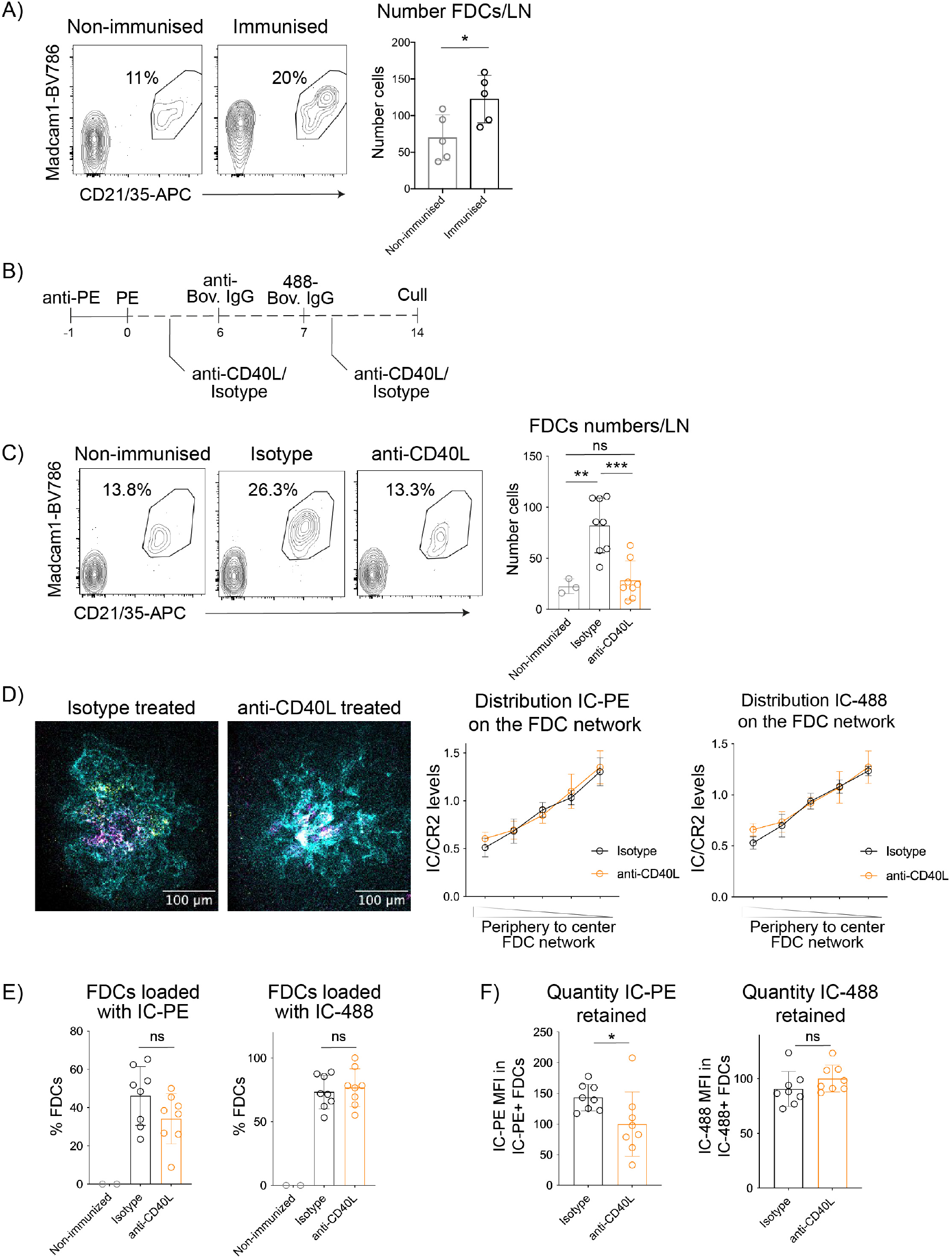
GC-derived signals induce FDC network expansion and activation but do not regulate antigen centralization or retention. A) Flow cytometry plots gated on FDCs as shown in Fig S1H and numbers of FDCs in LNs of non-immunized mice and mice immunized with IC-PE 13 days post-immunization (n = 5 mice). B) Experimental workflow to analyse the effect of CD40 signalling on antigen retention by FDCs. Mice were immunized first with IC-PE and then treated with anti-CD40L or isotype control antibodies (200 μg/mouse; 2 subsequent days). Seven days after the first immunization, mice were injected with the second IC (IC-Bovine-488) and treated again with anti-CD40L or isotype control. Draining LNs were analyzed 7 days after the second immunization. C) Flow cytometry plots and numbers of FDCs in non-immunized mice or mice immunized and treated with either anti-CD40L antibody (orange) or isotype control (black) as described in (B) (n = 3-8 mice from 2 experiments). D) Confocal images and quantification of the ICs distribution in mice treated as described in (B). Left graph shows the distribution of the first antigen (14 days post injection), right graph shows the distribution of the second (7 days post injection) (n = 8 LNs from 4 mice). E) Quantification of the percentage of FDCs loaded with IC-PE (1^st^ IC, left graph) and IC4-88 (2^nd^ IC, right graph) in anti-CD40L or isotype control-treated mice as described in (B) (n = 3-8 mice from 2 experiments). F) Quantity of ICs retained by FDCs measured as MFI in mice treated with anti-CD40L or isotype control antibodies (n = 8 mice from 2 experiments). Quantitative data shows the mean ± SD. T-test and One-way ANOVA with multiple comparisons, ns, P > 0.05; *, P < 0.05; **, P < 0.01; ** *, P < 0.001

First, we analyzed antigen distribution in mice unable to undergo BCR-induced activation because they express BCRs specific for non-cognate antigens. We injected WT mice, B1-8f (B1-8fl Igκ-/-, specific for the hapten NP) or MD4 (specific for hen egg lysozyme) BCR-transgenic mice with IC-PE and analyzed their LNs. Compared to the WT, the BCR transgenic mice had very few FDCs that formed poorly organized networks deficient at IC capture already at 24 h post-immunization (Suppl. Figure 2A and B). Thus, the basal level of BCR-driven B cell activation is necessary to promote or sustain full FDC development.

Second, we prevented T cell-dependent B cell activation and GC formation by blocking CD40 signalling(Foy et al., 1994). We immunised mice with two subsequent fluorescent antigen-ICs on days 0 and 7 and treated them with anti-CD40L blocking antibody or isotype control after each immunization (Figure 2B). The anti-CD40L antibody completely blocked the GC reaction (Suppl. Figure 2C) and the expansion of FDCs (Figure 2C) induced upon immunization. But even though the FDC expansion was impeded, the quantification of the antigen distribution in clarified LNs from the two groups of mice showed a similar centralization on day 14 (Figure 2D). In addition, the anti-CD40L treated mice had a similar percentage of FDCs loaded with either the first or the second IC compared to the control group (Figure 2E), and their FDCs retained similar amounts of the second injected antigen-IC (7 days post-injection), while containing slightly less of the first IC (14 days post-injection) (Figure 2F).

These results indicate that CD40L-dependent signals promote FDC network expansion after immunization, but that the increase in FDC cellularity is not responsible for the centralization of the antigen. CD40L-driven signals are also not essential for antigen retention on central FDCs for 7 days, although they may contribute to antigen retention of the FDC network at later time-points.

### Two different LZ FDC populations with different phenotypic and functional characteristics orchestrate antigen retention

To explore the possible differences between the peripheral FDCs that retain ICs for only a few days and the central FDCs that retain ICs for several weeks, we took advantage of sequential immunizations on days 0 and 6 with two differently labelled antigens (Figure 3A). On day 7 post-immunization, the first antigen is located only on the central FDCs (7 days post-injection), while the second antigen (24h post-injection) is located on both the central and the peripheral FDCs. This allowed us to distinguish the central FDCs from the peripheral by flow cytometry as FDCs positive for both antigens versus just the second antigen, respectively (Figure 3B). Using this strategy, we analysed the surface expression of FDC markers. Central FDCs expressed higher levels of FCGR2B and FCER2A than peripheral FDCs (Figure 3C). Central FDCs also expressed higher levels of CR2 (Figure 3D) but similar levels of the stromal marker Podoplanin (Figure 3E). Similar observations were made when comparing FDCs that retained antigen at day 7 to those that did not (Suppl. Figure 3A). A subset of FDCs expressing higher levels of FCGR2B and FCER2A had been previously suggested to be an activated state of FDCs in the GC LZ, induced upon immunisation(Allen and Cyster, 2008; Yoshida et al., 1993). Recently, single-cell transcriptomics revealed two subsets of FDCs present in the follicles at the steady state. LZ FDCs are located in a peripheral area closer to the subcapsular sinus, part of which becomes the LZ of the GC after immunization, while DZ FDCs, are located closer to the T cell zone(Pikor et al., 2020; Rodda et al., 2015). To understand if the central and peripheral FDC populations relate to the previously reported FDC populations, we analysed the expression of myosin heavy chain 11 (MYH11), a marker of the LZ FDC population (Suppl. Figure 3B)(Pikor et al., 2020). Both central and peripheral FDCs expressed high levels of MYH11, indicating that both subsets are LZ FDCs.

**Figure 3.**
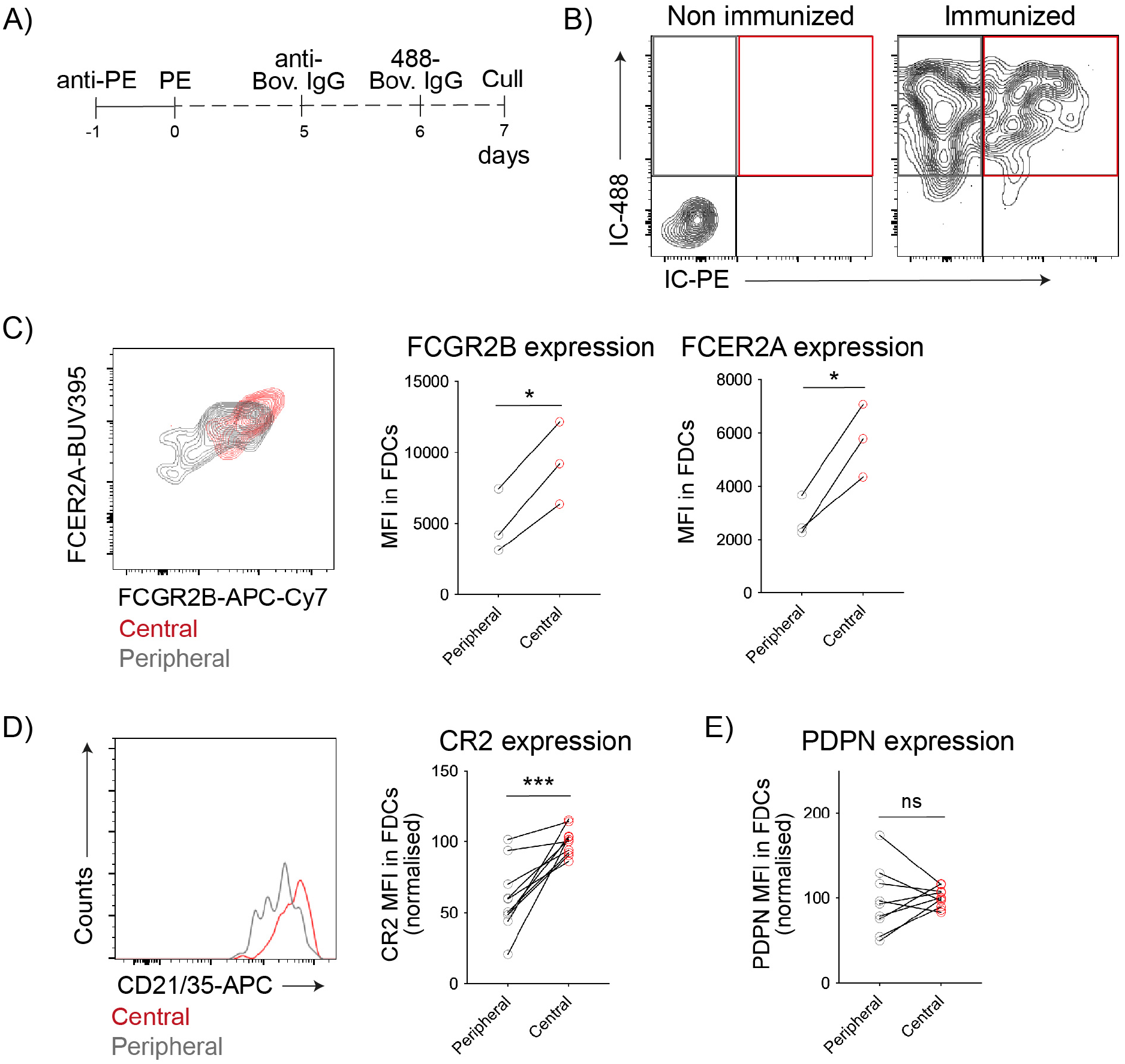
Central and peripheral LZ FDCs express different levels of IC-binding receptors. A) Immunization workflow to distinguish peripheral and central FDCs by flow cytometry. Mice were immunized first with IC-PE, followed by IC-488 6 days later. LN were analyzed 24 h after the second immunization. B) Flow cytometry gating strategy to identify peripheral and central FDCs in LNs of immunized mice following workflow described in (A). Retention of the first IC (IC-PE) distinguishes central (IC-PE^+^ IC-488^+^, red gate) from peripheral FDCs (IC-PE^-^ IC-488^+^, grey gate). C) FCGR2B (stained with anti-CD16/32 antibodies) and FCER2A (stained with anti-CD23 antibodies) membrane expression on central (red) and peripheral (grey) FDCs from mice immunized as described in (A) and (B) (n = 3 mice). D) CR2 membrane expression on FDCs as described in (A) and (B) (n = 10 mice, 3 experiments). E) PDPN membrane expression on FDCs as described in (A) and (B) (n = 10 mice, 3 experiments). Paired t-test, ns, P > 0.05; *, P < 0.05; ** *, P < 0.0005.

To understand better the relation between the peripheral and central LZ FDCs, we performed scRNAseq of the B follicle reticular cells marked by CXCL13-cre TdTomato(Pikor et al., 2020). Mice were analysed at two different time-points after immunisation with antigen-IC (Suppl. Figure 4A). Surprisingly, we did not observe major differences in gene expression at the different time points, therefore we performed all subsequent analyses on pooled samples. Unsupervised analysis revealed 10 distinct clusters of cells. Two of them corresponded to contaminating hematopoietic cells identified by their lack of *Cxcl13* expression and high expression of *H2-Aa* (Suppl. Figure 4B). The other 8 clusters corresponded to follicular stromal cells of which 7 were similar to clusters previously described using a similar experimental approach(Pikor et al., 2020; Rodda et al., 2018) (Figure 4A). Assignment of these 7 clusters using subset-specific genes identified two clusters of FDCs sharing *Cr2* expression; MRCs expressing *Madcam1* and *Tnfsf11*; IFRCs showing *Tnfsf11* and *Ch25h* expression; TBRCs with the expression of *Fmod* and *Ccl21a*; and two clusters of MedRCs sharing *Nr4a1* expression. The eighth cluster corresponded to follicular stromal cells with a prominent interferon-related signature, which we speculate could correspond to an activated stromal subset.

**Figure 4.**
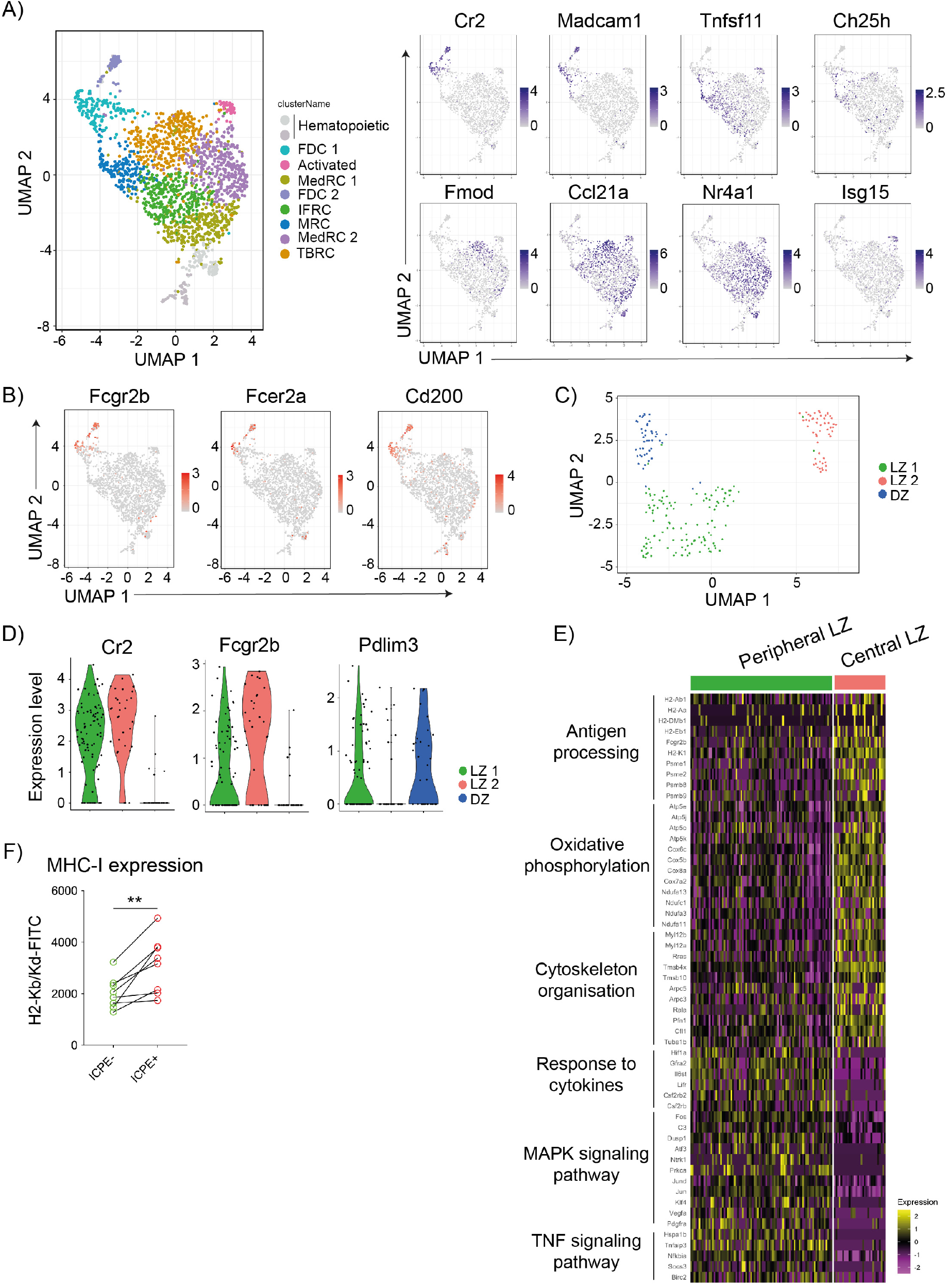
Single-cell transcriptomics differentiates two LZ FDC clusters with different functional activity. A) UMAP of Cxcl13-TdTomato expressing LN stromal cells. Left panel shows the merged data from mice immunized as described in Suppl. Figure 4A. Right panels show feature plots illustrating the expression of subset-specific marker genes for the indicated LN stomal cell clusters. B) Feature plots showing the expression pattern of Fcgr2b, Fcer2a and Cd200. C) UMAP of FDC1 and FDC2 cells from (A) after subclustering. D) Violin plots showing the expression of marker genes specific for the three FDC clusters (LZ 1 in green, LZ 2 in red and DZ in blue). E) Heatmap of scaled expression of genes differentially expressed between LZ 1 and LZ 2 FDC subsets grouped into functional and signalling pathways identified by STRING. F) MHC-I surface levels on IC-PE^+^ FDCs (central LZ FDCs) or IC-PE^−^ (peripheral FDCs) 7 days after immunization (n = 9 mice, 2 experiments). Paired t-test, *, P < 0.05.

Further analysis of the two FDC clusters indicated heterogeneity within the FDC 1 cluster. For example, only a fraction of cells in FDC cluster 1 expressed *Fcgr2b, Fcer2a* and *Cd200*, while these genes were expressed homogenously in FDC cluster 2 (Figure 4B). Furthermore, the cells in FDC cluster 1 that did not express these genes expressed higher levels of *Cxcl12* (Suppl. Figure 4B). Consequently, we decided to re-embed FDC 1 and 2 cells and perform a subclustering analysis, which grouped the cells into three FDC clusters (Figure 4C). Within these clusters, we were able to assign one cluster to DZ FDCs because of high *Pdlim3* and low *Cr2* expression, and two different clusters of LZ FDCs based on their high *Cr2* and *Myh11* expression (LZ 1 and LZ 2) (Figure 4D and Suppl. Figure 4C). LZ 1 showed lower *Fcgr2b* and *Fcer2a* expression relating this population to the peripheral FDCs described by flow cytometry, while LZ 2 cluster expressed higher levels of *Fcgr2b* and *Fcer2a*, relating them to the central FDC (Suppl. Figure 4C, Figure 4D, 3C). To confirm the relationship of the LZ 1 and LZ 2 clusters to peripheral and central FDCs in situ, we stained clarified LN 7 days after immunization with IC-PE with antibodies specific for FCGR2B. These images showed that central FDCs that retained antigen colocalised with the brightest staining for FCRG2B, indicating that the LZ 2 cluster corresponds to central FDCs (Suppl. Figure 4D).

To explore additional differences between the central and peripheral LZ FDC subsets, we analysed pathways enriched among their differentially expressed genes (DEG) using STRING(Szklarczyk et al., 2019) (Figure 4E). Central LZ 2 FDCs showed upregulated expression of molecules related to antigen presentation to B cells, but also to T cells, such as MHC-I and MHC-II related genes, together with several subunits of the proteasome complex. An increase in MHC-I expression on central versus peripheral FDCs was also observed at the protein level by flow cytometry (Figure 4F). Central LZ FDC also showed upregulated expression of genes involved in the mitochondrial respiratory complex participating in oxidative phosphorylation. We also observed that genes controlling cytoskeleton organisation were differentially expressed between central and peripheral LZ FDCs, which could explain the high degree of dendritic organisation and compaction that central FDCs show by microscopy. On the other hand, peripheral LZ FDCs seemed to be more responsive to extracellular signals with the upregulation of different cytokine receptor genes and intermediaries of MAPK and TNF signalling pathways (Figure 4E).

These results suggest that two different populations exist within LZ FDCs and that they likely correspond to the central and peripheral FDCs identified by imaging. Transcriptomics supports the specialized functionality of the two subsets, with an enhanced antigen-presentation role of the central LZ FDCs.

### Central and peripheral LZ FDCs show similarly low degradation of antigens

To understand the mechanisms underlying the long-term retention of antigens exclusively by the central LZ FDCs, we compared the ability of peripheral and central LZ FDCs to keep antigens in their native conformation. FDCs have been described to recycle antigens in non-degradative compartments, promoting the display of antigens for long periods of time compared to more degradative cellular subsets (e.g. DCs) also participating in antigen presentation to B cells(Batista and Harwood, 2009; Heesters et al., 2013). To study if the peripheral loss of antigen in the network could be due to increased antigen degradation by the peripheral LZ FDCs, we generated an antigen-degradation sensor based on labelling an antigen (bovine-IgG) with Atto488 and its quencher (BHQ-1), along with a non-quenchable dye AF647 (Figure 5A). BHQ-1 absorbed the spectral emission of Atto488 when the antigen was intact, but the fluorescence was recovered once the antigen was proteolyzed (Suppl. Figure 5A). To analyse the functionality of the antigen-quencher sensor in measuring physiological levels of antigen degradation, we incubated naïve primary B cells with 1μm beads coated with anti-IgM + fluorescent antigen (Bovine IgG-AF647-Atto488) or fluorescent antigen-quencher (Bovine IgG-AF647-Atto488-BHQ-1) at different time-points. Primary B cells can phagocytose antigen-coated particles, rapidly process the antigens and present antigen-derived peptides to CD4 T cells(Martínez-Riaño et al., 2018). We observed that the percentage of B cells containing quenched antigen (Atto488 low+ cells) decreased over time when the cells had been incubated with beads containing the antigen-quencher, but not the antigen alone (Suppl. Figure 5B). Similarly, the total antigen degradation levels, measured as a ratio between Atto488 and AF647 fluorescence, increased after 3 and 6 hours of incubation. Thus, this sensor detects physiological levels of antigen degradation.

**Figure 5.**
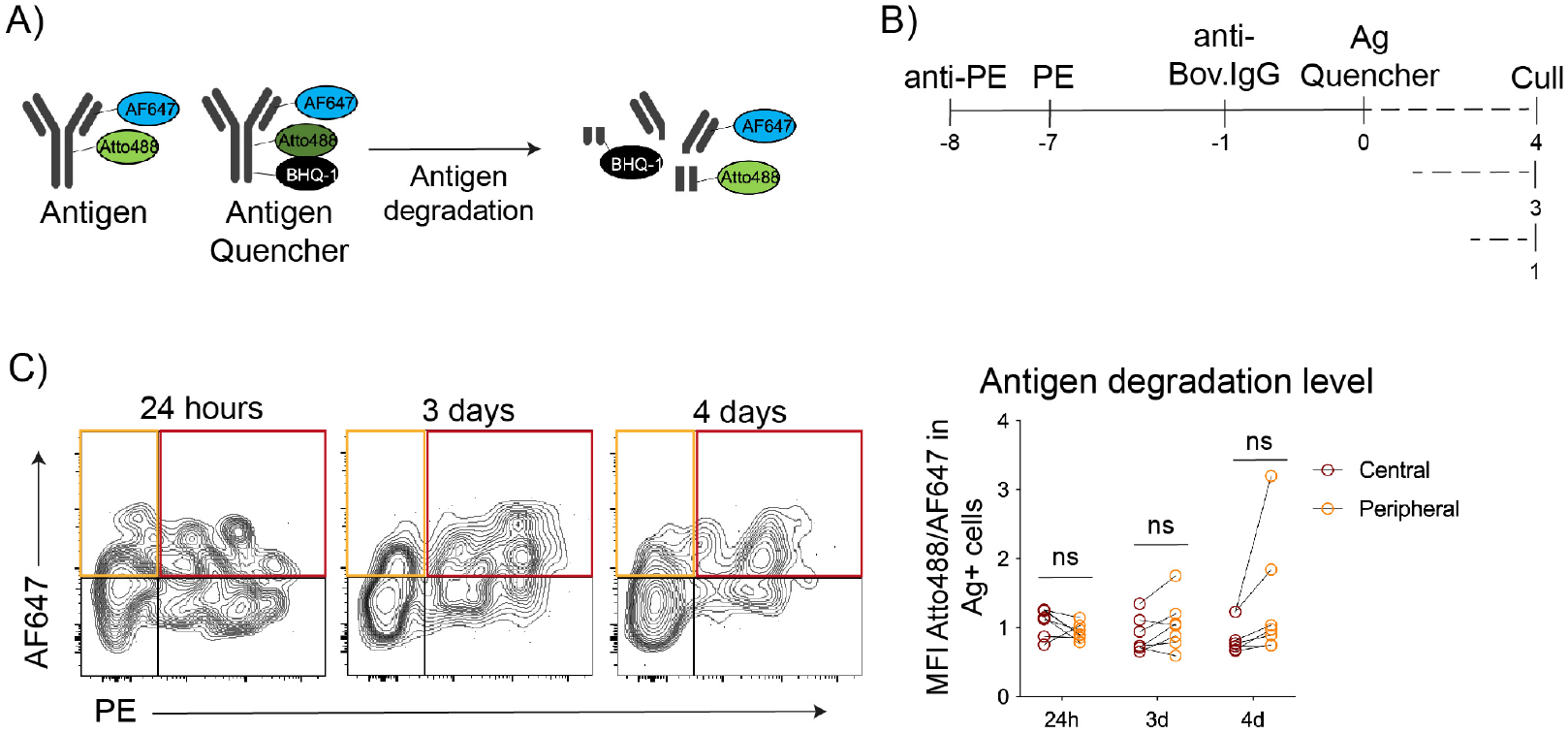
Central and peripheral LZ FDCs show a similarly low ability to degrade antigens. A) Schematics of the antigen-sensor and a control sensor lacking the quencher. The antigen (bovine-IgG) is covalently bound to AF647 and Atto488 dyes. The coupled BHQ-1 quencher absorbs Atto488 emission. Upon degradation, BHQ-1 is separated from Atto488, releasing its fluorescence from quenching. B) Immunization workflow to test the level of antigen degradation by central and peripheral LZ FDCs 24-hours, 3- and 4-days after immunization with the IC-antigen-degradation sensor. C) Left, flow cytometry gating on central (red) and peripheral (orange) FDC populations. Right, the levels of antigen-sensor degradation on central (IC-PE^+^ IC-antigen sensor^+^FDCs; in red) and peripheral (IC-PE-IC-antigen-sensor^+^ FDCs; in orange) FDCs in mice immunized as described in (B) (n= 7 mice, 2 experiments). Quantitative data shows the mean ± SD. Paired t-test, ns, P > 0.05.

To measure antigen degradation by different FDC subsets in vivo, mice were first immunised with IC-PE to differentiate central and peripheral LZ FDCs and, 7 days later, immunized with the antigen-quencher-IC. LN cells were analyzed at 24 hours, 3- and 4-days (Figure 5B). As expected, FDCs showed lower antigen degradation compared to B cells (Suppl. Figure 5 C and D). However, a comparison of the IC-PE-positive central and the IC-PE-negative peripheral FDCs showed similar sensor degradation (Figure 5C), indicating that the loss of antigen in the follicle periphery is not caused by increased antigen degradation in this area.

### Differences in CR2 density between central and peripheral LZ FDCs control antigen loss from the FDC network

One mechanism for enhanced antigen retention in the center of B cell follicles is a lower rate of dissociation of ICs from surfaces of central LZ FDCs compared to peripheral LZ FDCs. Since central LZ FDCs express higher levels of FCGR2B than peripheral LZ FDCs, they may bind IgG-containing ICs more stably. However, enhanced expression of FCGR2B on FDCs after immunization required CD40L-induced signaling, similarly to the expression of other activation makers such as FCER2A (Suppl. Figure 6A). Given that the retention of ICs in the follicle center was not affected by CD40L blockade (Figure 2E-G), it is unlikely that enhanced levels of FCGR2B on central LZ FDCs are responsible for selective antigen retention on these cells.

In contrast, surface levels of CR2 on FDCs, which were also higher on central LZ FDCs compared to peripheral LZ FDCs (Figure 3D, Suppl. Figure 3A), were independent of immunization or CD40L blockade (Suppl. Figure 6A). To understand if CR2 expression was different between central and peripheral FDCs at the steady state, we quantified its expression in clarified LNs from non-immunised mice by microscopy and normalised it to PDPN (Figure 6A), which was similarly expressed on all FDCs (Figure 3E). We observed that the normalized expression of CR2 was higher towards the center of the follicle, indicating that steady-state central LZ FDCs express more CR2 than peripheral LZ FDCs (Figure 6A).

**Figure 6.**
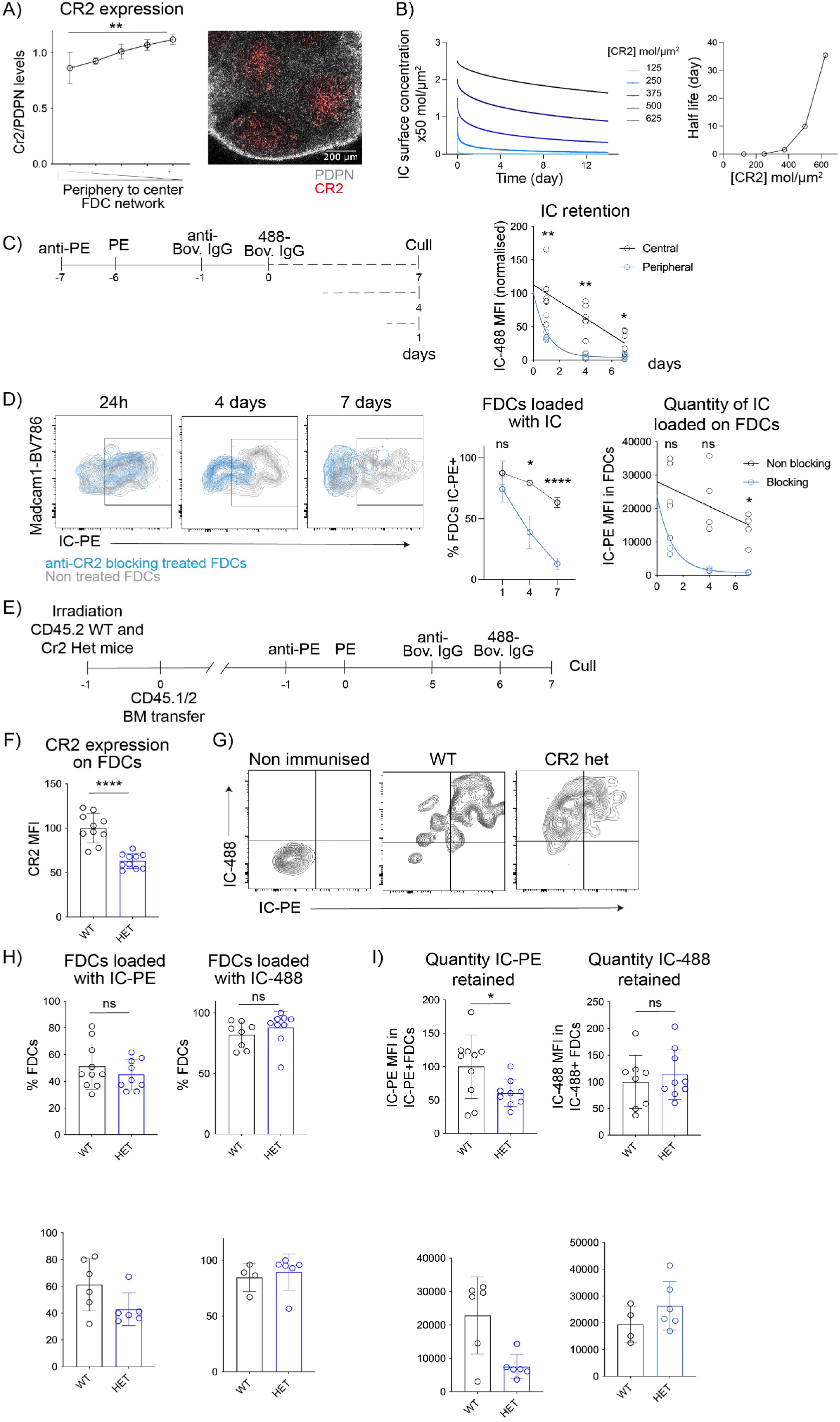
Membrane levels of CR2 dictate the half-life of antigen retention on peripheral and central FDCs. A) Quantification of CR2 levels from the periphery to the center of the FDC network in non-immunized mice. Anti-CR2 staining is normalised to that of PDPN (n = 4 LNs from 4 mice). Confocal image shows PDPN (grey) and CR2 (red) expression in a non-immunized mouse LN. B) Modelling IC dissociation from FDC surfaces. Left, surface concentrations of ICs retained on FDCs over time depending on CR2 expression levels (125 to 625 molecules/μm^2^). Right, plot of IC half-life versus CR2 surface concentration (molecules/μm^2^). A full description of the model is in Methods and parameters are illustrated in Suppl. Figure 6H. C) The rates of IC loss from central and peripheral LZ FDCs. Left, immunization workflow. 7 days after the first IC injection (IC-PE), mice were injected with a second IC (IC-488) and analyzed 24 hours, 4- and 7-days later. Graphs shows the quantity of the second IC (IC-488 MFI) retained by the central FDCs (IC-PE^+^ IC-488^+^, in grey) and peripheral FDCs (IC-PE^-^ IC-488^+^, in blue) (n = 7 mice, 2 experiments). Lines show non-linear regression fits to the data. D) Mice immunized with IC-PE were treated or not with 2 μg per site of anti-CD21/35 blocking antibody as described in Suppl. Figure 6J and analyzed 24 hours, 4- or 7-days later. Flow cytometry plots show IC-PE levels on gated FDCs from the two groups of mice. Graphs show the quantification of the percentage of IC-PE^+^ FDCs and the quantity of IC-PE (IC-PE MFI) on the total FDCs (n = 4 mice). E) Experimental workflow for measuring antigen retention by Cr2^+/+^ and Cr2^+/-^ FDCs. Lethally irradiated CD45.2 Cr2^+/+^ (WT) and Cr2^+/-^ (HET) mice were reconstituted with bone marrow from WT CD45.1/CD45.2 mice and 6 weeks later immunized first with IC-PE and then 7 days later with IC-488. LN were analyzed 24h after the second immunization. F) CR2 membrane expression on FDCs from WT (black) and Cr2 HET (blue) mice reconstituted as described in (E) (n = 10 mice). G) Gating strategy to analyse the IC loading in non-immunized WT mice, and WT or Cr2 HET mice immunized as described in (E). H) Percentage of FDCs loaded with IC-PE and IC-488 after 7 days and 24h postimmunization as described in (E) (n = 8-10 mice). I) Quantity of IC loaded (MFI) on total FDCs from WT and Cr2 HET mice following the experimental workflow described in (E) (n = 8-10 mice). Quantitative data shows the mean ± SD. Unpaired t-test and Two-way ANOVA, ns, P > 0.05; *, P < 0.05; **, P < 0.01; ** *, P < 0.001; ** **, P < 0.0001.

To validate that the retention of antigen-ICs by FDCs was CR2-mediated, lethally irradiated CD45.2^+^ WT and *Cr2* KO mice were bone-marrow reconstituted with CD45.1/2 WT haematopoietic cells and immunised with 2 subsequent antigen-ICs (Suppl. Figure 6B). Since stromal cells are radioresistant, bone marrow transplantation restricts the *Cr2*-deficiency specifically to the FDC compartment(Endres et al., 1999; Kapasi et al., 1994). To analyse the antigen binding to *Cr2*-deficient FDCs, we modified our flow cytometry gating strategy and identified FDCs based on the expression of FCGR2B and VCAM1 (Suppl. Figure 6C). FDCs from *Cr2* KO mice didn’t express CR2 (Suppl. Figure 6D) and showed negligible IC binding at 24 h or 7 days post-immunization (Suppl. Figure 6E and F). Thus, the antigen-IC retention observed in this immunization model was CR2-dependent.

To understand if the ~1.5 fold difference in CR2 levels on the cell surfaces of peripheral versus central LZ FDCs could be responsible for different rates of antigen loss from these cells over time, we generated a stochastic framework to model the probability of IC survival on FDC surface over time. As inputs for the modelling, we used FDC CR2 surface densities determined by quantitative flow cytometry (Suppl. Figure 6G), and CR2 dissociation (K_OFF_) and association (K_ON_) rates with C3dg, a C3 fragment closely resembling C3d, determined by Biolayer Interferometry and scaled for the size and flexibility of CR2, along with an estimated number of available C3d binding sites per IC (N_L_) (Suppl. Figure 6H). After an initial short period of loading of FDCs with the IC, the model calculated the dissociation of the ICs from the FDC surfaces assuming no additional formation of ICs and an irreversible loss of ICs after dissociation. The model predicted that CR2 levels promote the initial loading of ICs onto FDCs (Figure 6B, left). More strikingly, however, the model indicated that increasing CR2 levels non-linearly reduce IC dissociation over the next ~14 days. For example, a 1.5-fold increase in CR2 density, from 250 to 375 molecules per μm^2^, similar to the difference between peripheral and central FDCs, led to an increase in the apparent half-life of the IC on the FDC surface from 1.2 hours to 1.5 days. This corresponded to 6.7 higher levels of IC on the FDCs on day 14. Plotting the inferred IC half-lives confirmed their dramatic sensitivity to CR2 density with the inflexion point around their physiological levels on FDCs (Figure 6B, right). Thus, even subtle differences in CR2 surface levels on FDCs can have a dramatic impact on IC dissociation at time scales relevant for the GC reaction.

To quantify the rates of IC loss from central and peripheral LZ FDCs after immunization in vivo, we measured antigen levels on FDCs from mice immunized first with IC-PE, then with IC-488 7 days later, and sacrificed 1, 4 or 7 days afterwards (Figure 6C). The presence of the first IC allowed us to distinguish central (IC-PE^+^) from peripheral (IC-PE^-^) LZ FDCs and measure the quantity of IC-488 retained by each subset. The quantity of antigen was lower on the peripheral LZ FDCs on day 1 and dropped faster thereafter as compared to the central cells, as the model predicted. Thus, the rates of loss of ICs from peripheral and central FDCs are compatible with the model of differential IC dissociation.

To test the role of CR2-density in antigen retention, we used two strategies. First, we used an anti-CD21/35 blocking antibody to decrease the levels of CR2 on FDC and B cell surfaces available for binding of C3d. We titrated the anti-CD21/35 antibody to block approximately 50% of CR2 on FDCs, thus reducing the levels of CR2 on central LZ FDCs to the levels observed on the peripheral LZ FDCs in untreated mice (Suppl. Figure 6I). Mice were immunised with IC-PE for 24h, 4- and 7-days and treated with blocking anti-CD21/35 antibodies every two days until day 3 post-immunisation (Suppl. Figure 6J). The percentage of FDCs loaded with IC-PE, and the antigen quantity decayed faster in the anti-CD21/35 treated mice (Figure 6D), following similar kinetics to the one observed on peripheral LZ FDCs in untreated mice (Figure 6C). Second, we manipulated CR2 levels selectively on FDCs, by reconstituting lethally irradiated CD45.2 WT and *Cr2*-heterozygous (HET) mice with WT CD45.1/2 bone marrow. Reconstituted mice were immunized 6 weeks later with two subsequent antigen-ICs as above (Figure 6E). Radioresistant FDCs from *Cr2* HET mice expressed approximately 60% of CR2 on their membranes as did WT FDCs (Figure 6F). We observed that the percentage of FDCs loaded with the first and the second IC was not significantly different between both groups of mice (Figure 6G and H). However, we detected a significant reduction in the quantity of IC-PE displayed by *Cr2*-HET FDCs 7 days after immunisation, which was not observed at early time-points (IC-488) (Figure 6I). Thus, *Cr2*-HET FDCs had a similar ability to capture, but an increased loss of ICs within 7 days after immunization as compared to WT.

Collectively, these data indicate that the small differences in the level of CR2 expressed by the two subpopulations of LZ FDCs at the steady state result in preferential retention of antigens in the center of the follicle over time because of slower dissociation of ICs from central LZ FDCs.

### Retention of antigens in repeated dosing is driven by dynamic competition for central LZ FDCs leading to partial saturation and partial replacement

Having observed that the long-term retention of antigens takes place exclusively in the center of the FDC network, we wondered if central LZ FDCs could get saturated and become unreceptive to new antigens after repeated immunizations. We immunized mice consecutively with three (IC3; Figure 7A) or four (IC4; Figure 7B) antigen-ICs marked with different labels in one-week intervals. We analysed the FDCs according to the combination of antigens that they contained 24 h after the last immunization (Suppl. Figure 7A). FDCs that retained all the previous antigens showed the highest CR2 expression (Figure 7A and B, left panels), suggesting that they correspond to the central LZ FDCs. Indeed, imaging showed that all antigens older than a week were localised in the center of the follicle (Figure 7C). In the three-dose regime, these central FDCs containing all antigens (IC-647, IC-488 and IC-PE) were found to be loaded with more of the last antigen (IC-PE) than the FDCs containing only two, the second and the third antigens (IC-488 and IC-PE), or one, the third antigen (IC-PE) (Figure 7A right panel), which corresponds to our previous finding that peripheral FDCs hold less antigen than central FDCs 1 day after immunization. A similar phenomenon was detected in the four-dose regime, but only up to the third antigen (FDCs containing IC-647, IC-488 and IC-PE) (Figure 7B middle panel). In contrast, in the four dose-regime, the central FDCs containing all four antigens (IC-405, IC-647, IC-488 and IC-PE) captured similar amounts of the fourth antigen as the peripheral FDCs that contained only the last one (IC-PE). These data suggest that the central LZ FDCs captured the last antigen less efficiently than the previous ones, suggesting that these cells get saturated after three consecutive antigen-IC immunizations, although only partially.

**Figure 7.**
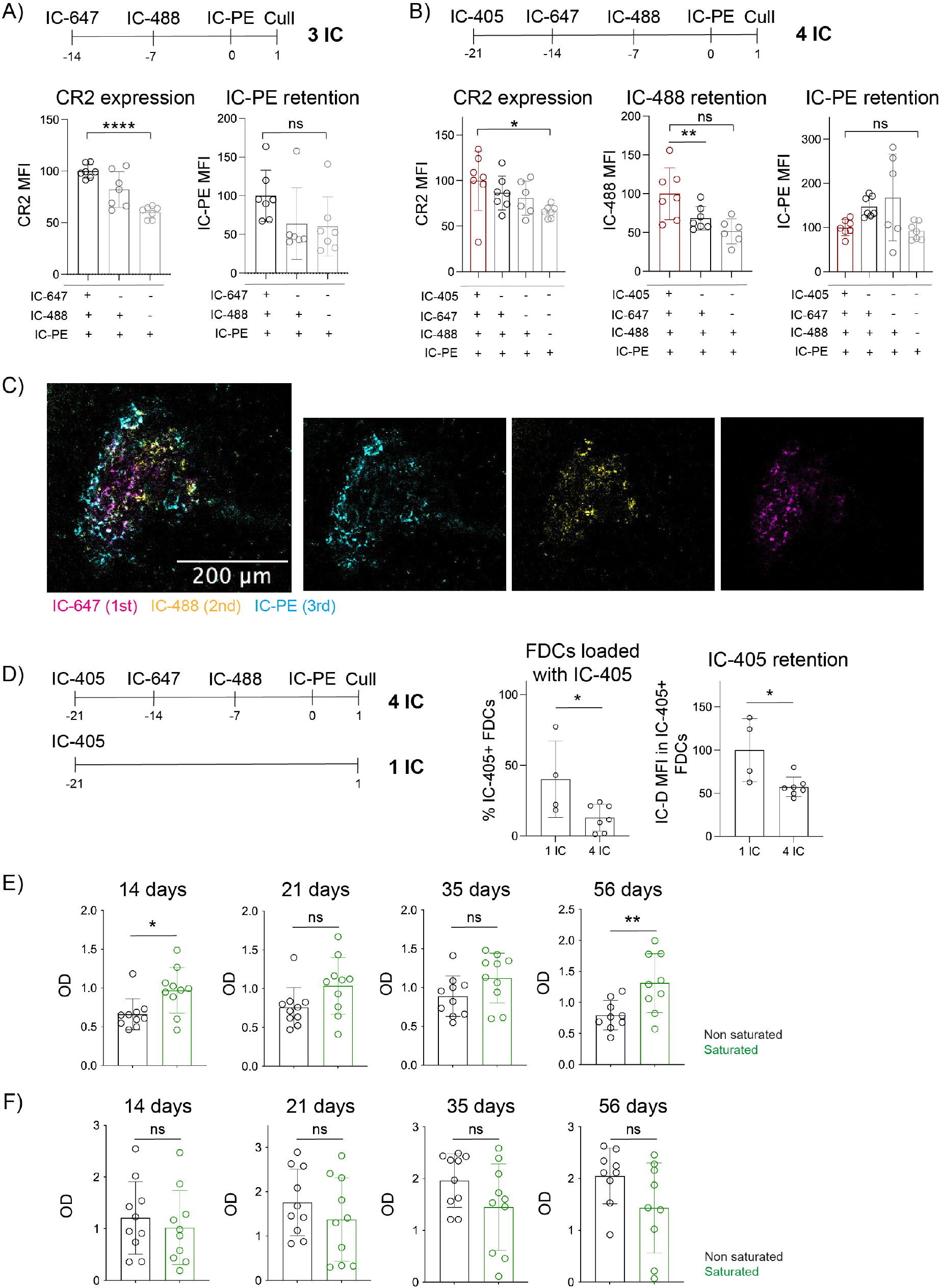
Successive immunizations partially saturate central FDCs, partially displace previous antigens, but still generate an efficient B cell response. A) Schematic workflow of mice immunized with three consecutive ICs containing distinct antigens. Plots show CR2 expression and retention of the last injected IC (IC-PE) in the different FDC subsets identified by the presence of all the three injected ICs (IC-647, IC-488 and IC-PE; black), the last 2 ICs (IC-488 and IC-PE; dark grey) or only the last one (IC-PE; light grey). FACS gating strategy used to differentiate the different FDC subsets is described in Suppl. Fig 7A. (n = 7 mice) B) Schematic workflow of mice injected consecutively with 4 different fluorescent antigen-ICs. Plots show CR2 expression and retention of the penultimate IC (IC-488) or the last IC (IC-PE) in the FDC subsets containing four injected ICs (IC-405, IC-647, IC-488 and IC-PE; red), the last three ICs (IC-647, IC-488 and IC-PE; black), the last two ICs (IC-488 and IC-PE; dark grey) or only the last IC (IC-PE; light grey). FACS gating strategy used to differentiate the different FDC subsets is described in Suppl. Fig 7A. (n = 7 mice) C) Confocal image of an FDC network from a LN of a mouse immunized with 3 different fluorescent antigen-ICs as described in (A). D) Experimental workflow to analyse the antigen displacement by subsequent immunizations. Mice were immunized with IC-405 alone or followed by four subsequent immunizations with different ICs. Graphs show the percentage of FDCs loaded with the first antigen and the quantity (MFI) of the loaded antigen in the two groups of mice (n = 7 mice). E) Levels of NP-specific IgM in sera of mice 14, 21, 35 and 56 days after immunization with either two antigen-IC regime (No saturation condition; black) or four antigen-IC regime (Saturation condition; green) as indicated in Suppl. Figure 7B. (n = 9-10 mice) F) Levels of NP-specific IgG1 in sera of mice immunized as described in (E). (n = 9-10 mice) Quantitative data shows the mean ± SD. Unpaired t-test and One-way ANOVA, ns, P > 0.05; *, P < 0.05; **, P < 0.01; ** *, P < 0.001; ** **, P < 0.0001.

To understand if the converse also happens, that is that old antigens get displaced by immunization with new ones, we immunised mice with either one or four consecutive antigen-ICs (Figure 7D). We analyzed the FDCs 22 days after the first immunization (1 IC) or one day after the last one (4 IC), respectively. We observed that the percentage of FDCs loaded with the first antigen-IC (IC-405) and the amount of IC-405 on these FDCs was lower when the mice were immunized with the three additional immunizations than when they were not. This suggests that the dissociation of the oldest antigen from the FDCs was enhanced by competition with the new antigens, but that it was not completely replaced.

To understand if the partial saturation and partial replacement of antigens on central LZ FDCs could modify the B cell response to new antigens, we immunised mice with three subsequent antigen-ICs or only one antigen-IC and challenged them with IC-NP 21 days after the first immunization (Suppl. Figure 7B). We tracked the antibody response generated to NP in the two groups of mice over the next 56 days. We observed that the high-affinity NP-specific IgM (Figure 7E) and IgG1 class-switched responses (Figure 7F) were similar between both groups of mice, or even slightly higher in IgM antibody production under saturation conditions at the first and last time-points (Figure 7E). Thus, the partial saturation of the FDCs by the presence of previous antigens doesn’t impede the development of the B cell response to new, unrelated antigens.

## DISCUSSION

Our study describes the mechanisms controlling antigen dynamics on the FDC network. We show that pre-existing topological differences in the network control the location and duration of antigen retention. Specifically, two populations of LZ FDCs, peripheral and central, are responsible for the dynamics through different abilities to retain antigens over time. We found that the central LZ FDCs are selectively responsible for the long-term antigen display in the GC(Tew and Mandel, 1979) and function as an antigen reservoir containing multiple antigens from previous immunizations. In contrast, peripheral LZ FDCs only contain antigens in the early-time points after immunization. We implicate enhanced CR2 levels on central FDCs as the main factor in controlling antigen retention. This suggests that the avidity of antigen-ICs for FDC surfaces is the principal mechanism that controls the length of antigen availability to GC B cells, and the rules for FDC saturation and antigen replacement upon secondary immunizations.

The distinction between peripheral and central FDCs builds on the previously recognized heterogeneity of the follicular stroma and FDCs in particular. The central FDCs likely correspond to FDCs previously described as activated FDCs located in the LZ of the GC based on the expression of FCGR2B and FCER2A. FDC activation is likely a consequence of upregulated LTA1B2 production by GC B cells(Ansel et al., 2000). However, we show that peripheral and central FDCs can be distinguished even before immunization.

The mechanisms driving the distinction between the peripheral and central FDCs thus remain unclear. Basal production of LTA1B2 and TNF by B cells at the steady-state is necessary for the FDC development and maintenance(Endres et al., 1999; Fu et al., 1998; Gonzalez et al., 1998; Pasparakis et al., 1996). Unexpectedly, we observed that monoclonal BCR Tg mice have a defect in FDC formation, which could indicate that a low level of B cell activation at the steady-state, possibly induced by endogenous antigens(Noviski et al., 2019; Zikherman et al., 2012), contributes to FDC maturation or maintenance. Signals from GC B cells may thus promote the spatial heterogeneity of the FDC network. In addition to FDC activation, the FDC network also expands in response to immunization, likely due to the differentiation of new FDCs from subcapsular perivascular precursors(Jarjour et al., 2014; Prados et al., 2021). Speculatively, new FDCs arising at the subcapsular sinus may mature through peripheral FDCs into central ones. However, acute blockade of FDC network expansion using anti-CD40L antibodies did not alter the peripheral to central FDC ratio or the centralization of the antigens after immunization. It is possible, however, that FDC network remodelling may contribute to the topological heterogeneity at later time points when the newly differentiated FDCs integrate into existing networks or replace them. Understanding the cellular dynamics underlying FDC heterogeneity and antigen centralization to the GC may reveal additional mechanisms through which the stroma regulates the B cell response.

We offer further insight into the function of the central and peripheral LZ FDCs through single-cell transcriptomics. The transcriptomics data support a specialised role for a cellular cluster expressing markers of the central LZ FDCs in presenting antigens to B cells. Surprisingly, this subset also shows higher expression of genes involved in the processing and presentation of antigen-peptides (H2-Ab1, H2-Aa, H2-DMb1, H2-Eb1, H2-K1 and several subunits of the immunoproteosome), suggesting that they can actively process and present antigens to CD8 and CD4 T cells in a TCR-specific fashion, although the relevance of this needs to be further investigated. This data agrees with a recent report that described the expression of MHC-II and PD1L by human FDCs(Heesters et al., 2021). Transcriptomics also revealed differences in the metabolism of LZ FDCs subsets. The subset corresponding to central FDCs upregulates the expression of genes encoding Complex I, IV and V of the mitochondrial electron chain, suggesting a possible increase in oxidative phosphorylation. The central FDC subset also has a preferential expression of genes involved in the organization of the cytoskeleton which could suggest different membrane stiffness, contributing to a more stringent B cell affinity discrimination(Spillane and Tolar, 2017). Peripheral LZ FDCs instead seem to be more sensitive to the environment, expressing genes related to receptor and cytokine response and regulation of TNF and MAPK signalling.

One mechanism that could selectively prolong antigen retention by central FDCs is the internalization of ICs into non-degradative endosomal vesicles that recirculate to the surface(Heesters et al., 2013). This prevents dissociation and provides periodic exposure of the antigens to B cells, which has been suggested to impact the GC dynamics(Arulraj et al., 2021). However, we did not detect differences in surface versus intracellular antigen distribution between peripheral and central FDCs (not shown) and the in-vivo comparison of the antigen degradation also did not reveal differences. This suggests that intracellular trafficking does not contribute to the enhanced antigen retention capacity of central FDCs.

Similarly, we did not find evidence that enhanced Fc-receptor expression by central FDCs could promote antigen retention. Our data illustrate that upon IC immunization, almost all the antigen-IC binding observed on the FDCs is CR2 dependent, even after FCGR2B expression is upregulated. This is consistent with previous data showing FCGR2B to be dispensable for antigen capture during the primary response(Suzuki et al., 2009). However, some studies did find a role for FCGR2B upon reimmunization or under autoimmune conditions(Barrington et al., 2002; Poel et al., 2019; Qin et al., 2000). It is thus possible that FCGR2B expression has a regulatory role under some specific conditions.

Instead, we show that the different level of CR2 expression on the membrane of central and peripheral LZ FDCs is the major factor that drives the differences in their retention of antigens. The interactions of CR2 with C3d are low affinity and highly transient (K_D_ ~ 140 nM, half life ~ 6 s), mandating that any long-term retention involves avidity through multivalent binding of several CR2 molecules to multiple C3d fragments on the IC. This is consistent with the substantial length and flexibility of CR2 and with its expression level on FDCs, which is the highest in the body. Modelling indicates that the avidity is dependent on the surface density of CR2, with the predicted half-life of binding sharply rising when CR2 levels reach the physiologic range of expression on FDCs (>200 molecules/μm^2^). Central FDCs express approximately 1.5x more CR2, generating enhanced avidity for ICs and preventing IC dissociation, thus facilitating the retention of the ICs on central FDCs. This is also consistent with FDCs efficiently acquiring ICs in the first hours after immunization from CR2^+^ non-cognate B cells(Phan et al., 2007), which express only ~ 20 CR2 molecules/μm^2^.

The central role of IC avidity for FDC surfaces has implications for the principles of antigen retention in the GC after repeated challenges. We show that under repeating immunization, antigens compete for the CR2-FDC binding. For example, after 3 repeated IC doses, new antigens are not as efficiently loaded onto the FDCs. Conversely, we observed that older antigens located on the FDC network are displaced by new incoming antigens. This is consistent with a dynamic avidity binding model, where CR2 occupancy lowers CR2 availability and thus FDC avidity for the incoming new ICs and at the same time new ICs also lower the free CR2 accessibility, causing accelerated dissociation of previous ICs. This suggests that the FDC network is robust in the face of saturation and is able to capture enough antigens despite partial saturation to activate B cells and generate an efficient antibody response(Kosco-Vilbois, 2003). Indeed, we could not see differences in the antibody response under conditions of partial CR2 saturation versus non-saturation. This result contrasts with the idea of saturable FDC niches for GC development(Avancena et al., 2021). Thus, the dynamic avidity of antigens for two FDC populations spanning a range of CR2 levels may help to explain how the FDC network handles repeated incoming antigens, keeping a dynamic “memory” of past antigens but also remaining receptive to new ones.

In summary, we have shown two specialised populations of LZ FDCs with different abilities to retain and display antigens. The differential expression of CR2 between both populations drives the changes in antigen retention observed after immunization. Understanding the mechanisms that regulate the development of central and peripheral FDCs and their CR2 levels may help to determine their exact roles in supporting the B cell response. Loss of CR2 on FDC leads to contraction of the GC, and reduction of late antibody titres. Therefore, vaccination strategies could aim to promote a better complement deposition on vaccine materials to increase the avidity of the interaction with FDCs and sustain the availability of the antigens to B cells. Antigens with high complement fixation capacity may be more efficient in competition for CR2 on central FDCs. Immunization strategies taking advantage of this mechanism may prolong GC responses and enhance B cell affinity selection and the emergence of broadly neutralising B cell specificities(Klein et al., 2013; Pappas et al., 2014). Antigens with high CR2 avidity may also effectively displace previous antigens and modulate existing responses which may be useful in situations of chronically elevated IC loading, such as during systemic autoimmune diseases.

## ACKNOWLEDGEMENTS

This work was supported by the Francis Crick Institute, which receives its core funding from Cancer Research UK (FC001185), the UK Medical Research Council (FC001185) and the Wellcome Trust (FC001185). SW is supported by the National Science Foundation (NSF) Grant MCB-2225947 and an NSF CAREER Award PHY-2146581. We thank the Francis Crick Institute Animal facility, Light Microscopy, Flow Cytometry and Advanced Sequencing Technology Platforms. We thank Dinis Calado for the Cr2-KO (Cr2tm1Hmo) mice and Barton Haynes for the YU-gp120 pcDNA3.1 plasmid. We thank Laabiah Wasim for critical reading of the paper.

## AUTHORS CONTRIBUTION

A.M-R. designed and performed the experiments and analyzed the data. S.W. generated the IC-FDC mathematical dissociation model. S.B. analyzed the scRNAseq data. S.M. measured the CR2-C3dg binding rates. A.C. produced the YU-gp120-SpyTag protein. K.S. provided advice. B.L. provided the CXCL13-TdTomato mice and advice. P.T. designed the experiments and supervised the research. A.M-R. and P.T. prepared the manuscript.

## MATERIALS AND METHODS

### Mice

C57BL/6, *Cr2*-KO (*Cr2*^tm1Hmo^), CXCL13-TdTomato (Tg(*Cxcl13*-Cre/tdTomato)719Biat) and CD45.1 (B6.SJL-*Ptprc^a^ Pepc^b^* /BoyJ) mice were used. To obtain Cr2-HET and CD45.1/CD45.2 mice, Cr2-KO and CD45.1 mice were bred with C57BL/6 mice. All experiments were approved by the Francis Crick Institute and UCL Ethical Review Panels and the UK Home Office.

To generate bone marrow chimaeras, recipient mice were lethally irradiated with two doses of 5Gy and reconstituted with 5×10^6^ donor bone marrow cells by intravenous injection. Reconstituted mice were fed with 0.2 mg/ml Baytril (Enrofloxacin) in their drinking water for 4 weeks post-reconstitution.

### Immunization

Mice were immunized intraperitoneally with 200 μg anti-antigen antibody in 200 μL of PBS. 18h later, mice were injected sub-cutaneously with 10 μg of fluorescent antigen in 100 μL mixed 1:1 with Imject Alum Adjuvant (ThermoFisher) in the upper and lower flank to target the brachial, axillary and inguinal draining LNs. Mouse tissues were analyzed by flow cytometry, confocal microscopy and ELISA at different time-points after immunisation.

### Cellular isolation

For FDCs preparation, draining lymph nodes were disaggregated into small pieces with 25G needles and collected in RPMI-1640 medium containing 2% FCS, 20 mM HEPES pH 7.2, 0.1 mg/ml collagenase P (Roche) and 25 μg/ml DNase I (Sigma). Dissociated tissue was incubated at 37 °C for 60 min, recollecting supernatant every 15 min. After enzymatic digestion, cell suspensions were filtered using a 100 μm strainer and washed with PBS containing 0.5% FCS and 10 mM EDTA. Cell suspensions were used directly for staining with antibodies.

For B cell isolation, spleens were disaggregated using 40 μm strainer and treated with AcK buffer (made in house) for 5 minutes. Single-cell suspension was subsequently incubated with CD43 microbeads (Milteny Biotech), following manufacturer’s instructions.

### Staining

For flow cytometry staining, single-cell suspension was incubated with Fixable Viability Dye-e506 (Invitrogen) for 15 minutes in PBS. Cells were subsequently incubated with anti-CD16/32 (purified or labelled) and the appropriate antibodies for 20 minutes at 4 degrees in PBS containing 0.5% FCS and 10 mM EDTA for FDC analysis or PBS containing 2% FCS and 2mM EDTA for B cell analysis. The following antibodies were used for FDC and B cell phenotyping: B220 (RA3-6B2), CD45.2 (104), CD21/35 (7E9), PDPN (8.1.1), Madcam1 (MECA-367), CD31 (390), CD23 (B3B4), H2-Kb/Kd (28-8-6), CD95 (Jo2), IgD (11-23c), T-and B-Cell activation antigen (GL7), CD45.1 (A20). Cells were analysed using an LSR-Fortessa flow cytometer and analysed using FlowJo. For cell sorting experiments for scRNAseq, cells were sorted using FACS Aria Fusion and collected in PBS + 0.05% BSA.

For microscopy staining, LNs were incubated 4 hours with Antigenfix solution (Diapath) and washed and permeabilised in PBS containing 1% BSA, 1% normal mouse sera and 2% Triton X-100 for 24h. LNs were incubated with the antibody mix in the permeabilization buffer for 3 days at RT while shaking. Organs were subsequently washed in permeabilization buffer for 24h and incubated with RapiClear solution (1.47 RIN) for 24h at RT. Clarified organs were imaged mounted in RapiClear solution using Leica SP5 Upright or Leica SP8 Falcon Inverted microscopes. For PDPN staining, LNs were permeabilised in 4% SDS in 200mM boric acid at 37°C for 4h and labelled in 4% SDS solution for 4 days at RT as previously described in(Messal et al., 2021).

### Image analysis

LN image stacks were analyzed using MATLAB supplemented with the MIJ ImageJ plugin. Images were subsampled 4:1 in the x and y axes. The image channel containing anti-CD21 FDC staining was bandpass-filtered and thresholded to generate 3D masks identifying individual FDC networks. The FDC network masks were labelled and segmented into 6 concentric shells. The mean pixel intensity of the anti-CD21 and antigen fluorescence was then measured in each shell in each network. The data were background-subtracted and normalized by the mean pixel intensity of the corresponding signal of each follicle. To quantify antigen centralization, the normalized antigen intensity in each shell was divided by the corresponding normalized intensity of the anti-CD21 signal.

### Droplet-based single-cell RNA sequencing analysis

Sorted CXCL13-TdTomato^+^ PDPN^+^ live cells were run using the 10× Chromium (10× Genomics) system, and cDNA libraries were generated according to the manufacturer’s recommendations (Chromium Single-Cell 3’ Reagent Kit (v3.1 Chemistry)). Libraries were sequenced via Hiseq 4000 for Illumina sequencing. Raw sequencing data were processed using the CellRanger pipeline version 3 (10× Genomics) with the Ensembl GRCm38 release 89 reference transcriptome. Count tables were loaded into R and further processed using the Seurat R package version 3.1.5(Butler et al., 2018). Samples were pooled from three independent experiments (Cxcl13-Td immunized for 24h, LNs from six mice pooled; Cxcl13-Td immunized for 7 days with a first antigen-ICs and 24h with a second antigen-ICs, LNs from six mice pooled). Subsequently, cells containing fewer than 200 distinct genes and cells with more than 10% of unique molecular identifiers stemming from mitochondrial genes were excluded. Furthermore, cells that featured at least one read count for Lyve1, Hba-a1, Hba-a2, Krt18, Trac, Cd3d, Cldn5, Ly6c1, Egfl7, Ptprc, S100b, Cd79a, Cd79b genes were removed to eliminate contaminating hematopoietic cells, erythrocytes, endothelial and epithelial cells as previously described(Pikor et al., 2020). After quality control and removal of contaminants, the remaining cells were retained for further processing using the default method from the Seurat package (version 3.1.5). Clusters were characterised based on described markers(Pikor et al., 2020; Rodda et al., 2018). Differentially expressed genes between three FDC clusters were performed using Seurat. Pathways enriched among their differentially expressed genes (DEG) were analyzed using STRING(Szklarczyk et al., 2019).

### Protein labelling with dyes

Antibodies were conjugated to one of several fluorophores AlexaFluor 405, AlexaFluor 488, AlexaFluor 647 NHS esters (Thermo Fisher) in Sodium Carbonate buffer, according to the manufacturer’s instructions. Excess dye was removed using Zeba 7K MWCO desalting columns (Pierce, Thermo Fisher).

For the degradation sensor, antigen was conjugated first with AlexaFluor 647 and AlexaFluor 488 NHS esters (Thermo Fisher) and with BHQ-1 quencher as previously described in(Li et al., 2015).

### In vitro antigen degradation assay

Conjugated Bovine IgG with AlexaFluor 647 and AlexaFluor 488 NHS esters (control) and Bovine IgG conjugated with AlexaFluor 647 and AlexaFluor 488 NHS esters and BHQ-1 NHS ester (ratio 1 IgG: 9 BHQ-1 molecules) were treated at 50°C for 30 minutes and 95°C for 5 minutes with 2mg/mL Proteinase K. Fluorescent emission was measured using The Spark multimode plate reader (Tecan).

### Ex vivo degradation assay

50 × 10^6^ carboxylated latex beads 1 μm in diameter were incubated overnight with a concentration of 20 μg/ml of anti-IgM plus 20 μg/ml antigen-quencher or only anti-IgM in 1 ml of PBS at 4°C.

Naïve purified B cells were resuspended in complete RPMI (10% FBS, 100μM non-essential amino acids (ThermoFisher), 2mM L-Glutamine (ThermoFisher), 50μM 2-Mercaptoethanol (ThermoFisher) and Penicillin-Streptomycin (GE Healthcare Life Sciences)) and plated in 96-well V-bottom plates at a concentration of 0.5 × 10^6^ cells in 50 μl. Antibody-coated beads were added to reach a bead:cell ratio of 3:1. The cellular and bead suspension were briefly centrifuged at 400 *g* and were incubated at 37°C for different time points. Subsequently, cells were washed and stained on ice(Martínez-Riaño et al., 2018).

### HIV-multimeric nanoparticles

60-mer SpyCatcher-SpyTag particles were generated as described in(Bruun et al., 2018). Briefly, monomeric SpyCatcher-mi3 (kindly donated by Mark Howarth) was incubated with 3 times molar excess YU-gp120-SpyTag HIV envelope protein for 18h at 25°C in PBS and dialysed using 300 KDa MWCO membrane (Spectra/Por Float-A-Lyzer G2) in Sodium Carbonate buffer following manufacturer’s instructions. VLPs were subsequently incubated with AF-555 NHS ester for 1h at 25°C and dialysed using 300KDa MWCO membrane in PBS for 2 days. Mice were immunised with a dose containing 1 μg YU-gp120 protein in 100μL of PBS in the flanks.

### YU-gp120-SpyTag HIV envelope production

An SpyTag sequence was inserted in the N-terminal part of the YU-gp120 sequence. The recombinant protein was produced in 293F cells transfected with YU-gp120-Spytag expressing pcDNA3.1 plasmid (a kind gift from Barton Haynes) as described in(Liao et al., 2013). Briefly, cell supernatant was filtered with 0.8 μm filter, mixed with *Galanthus nivalis* lectin (GNL) binding buffer and load into a lectin-agarose column previously equilibrated with binding buffer (5 times the volume). After that, the column was washed 5 times with binding buffer and the protein was eluted using a Mannose solution. The purity was assessed by running an SDS-PAGE.

### CR2-C3dg binding measurement

The on- and off-rate and the equilibrium dissociation constant for the CR2 interaction with C3dg was measured using bio-layer interferometry (Octet, Sartorius). We loaded a his-tagged human CR2 protein (Bio-techne) to Nickel-NTA sensor at a concentration of 36.8 nM. Human C3dg protein was produced as described(Nagar et al., 1998). Briefly, human C3dg cDNA containing the Cys1010Ala mutation inserted into pET13b expression plasmid lacking the his-tag (a kind gift from J. Eisenman) was transformed into BL21 E. coli. After induction with IPTG, soluble C3dg was purified from bacterial lysates using CM Sepharose followed by gel filtration on a Superdex 200 column. Association of the C3dg protein with CR2 was measured for concentrations ranging from 0.023 μM to 2.9 μM. The equilibrium dissociation constant was determined by fitting the plateau values with a binding model, yielding K_D_ = 317 ± 30 nM. The association and dissociation rates were determined by fitting a kinetic model yielding K_ON_ = 616,154 M^−1^ s^−1^, K_OFF_ = 0.15 s^−1^.

### IC-FDC dissociation mathematical model

To describe the dynamics of IC dissociation from FDCs at varying CR2 concentrations, we generated a stochastic framework comprising microscopic events that alter the probability of IC survival over time. Specifically, as an IC is loaded onto the FDC membrane, a multiplicity of adhesive bonds forms between the C3d ligands coating the IC particle and the surface CR2. Individual dissociated CR2-C3d pairs can rebind, as long as some bonds remain to hold the IC. Once all bonds open, the IC is irreversibly lost.

Mathematically, starting from maximum bond formation between the IC and CR2 on the corresponding membrane patch (assuming that each membrane patch can host at most one IC), stochastic IC loss proceeds through a one-step master equation:

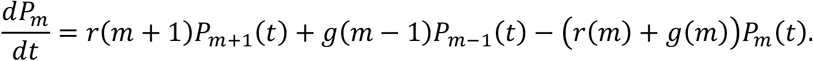

Here *P_m_*(*t*) represents the probability that *m* bonds remain closed between an IC and the FDC at time *t*, which evolves due to dissociation of any closed bond at an unbinding rate *r*(*m*) = *mk_off_* and formation of a new bond at a rebinding rate *g*(*m*) = *C*(*m*)*k_on_*. We set *g*(0) = 0 to avoid IC re-association. Rebinding shifts the equilibrium state away from complete dissociation, stabilizing multivalent binding (IC survival) in the presence of noise. Eventually, all bonds break if one waits long enough. The CR2-C3d single-bond dissociation rate, *k_off_*, was obtained from Biolayer Interferometry. In the model 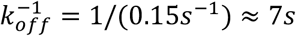 sets the time unit. *k_on_* was adjusted to recapitulate the typical half-life of ICs observed at known CR2 concentrations.

The key quantity in the rebinding rate, *C*(*m*), counts the number of binding configurations (bond arrangements) given *m* closed bonds. Importantly, the form of *C*(*m*) depends on bond properties (e.g. length and flexibility) and binding geometry (e.g. C3d spacing, curvature of the IC surface, distance of IC from the FDC surface when bound). We used the all-to-all binding scenario: *C*(*m*) = (*n_R_* – *m*)(*n_L_* – *m*), whereby each of *n_L_* C3d ligands on a guest IC is accessible to all n_R CR2 on the host membrane patch. This scenario is appropriate for long, flexible molecules like CR2. Note that *C*(*m*) increases rapidly with the valency *m* due to nonlinearity, resulting in a high sensitivity of IC survival to CR2 concentration.

We simulated the model and computed the time-dependent survival probability of an IC, 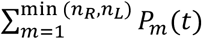, for a given CR2 concentration (*n_R_*). To account for variations of CR2 density for a given mean value (corresponding to different FDC populations), we consider a Poisson distribution of CR2 level among membrane patches for a certain mean CR2 density 〈*n_R_*〉: *P*(*n_R_*) = *e*^−〈*n_R_*〉)^〈*n_R_〉^*n_R_*^*/*n_R_*!. By averaging over this distribution, we obtain the mean IC survival rate 〈*S*(*t*)〉 and use 〈*S*(*t*)〉 · 〈*n_R_*〉 to represent the overall surface concentration of IC on the FDCs. In Figure 6B we plot this quantity against time and also extract the half-life (time taken to reach half of the initial IC level) at varying CR2 concentrations.

### anti-CD40L treated mice

Mice were immunised with a first antigen immunocomplex (Immunisation protocol) and five days later injected intraperitoneally with 200 μg anti-CD40L blocking antibody or its isotype control for two subsequent days. Mice were subsequently immunised with a second antigen immunocomplex and injected again two days later with 200 μg anti-CD40L blocking antibody or its isotype control for 2 subsequent days. Draining LNs were used for flow cytometry and microscopy.

### anti-CD21/35 blocking antibody treated mice

Mice were injected with 2 μg anti-CD21/35-BV421 (clone 7G6) subcutaneously in the upper and lower flank to target the brachial, axillary and inguinal draining LNs starting on the day of the experiment and every 2 days thereafter up to 3 injections. Mice were subsequently immunised with antigen-immunocomplexes (following the immunization protocol described above) and analysed after 24, 4 and 7 days post-immunisation.

### ELISA

In immunised mice, sera were obtained at 14, 21, 35 and 56 days after immunisation. Plate-bound NP(7)-BSA (10 μg/ml) was used to measure antigen-specific antibodies. Class-switched serum immunoglobulin levels were detected using SBA Clonotyping System HRP kit (Southern Biotech).

