## Supplementary Figures for "Long-term retention of antigens in germinal centres is controlled by the spatial organisation of the follicular dendritic cell network"

Supplementary Figure 1

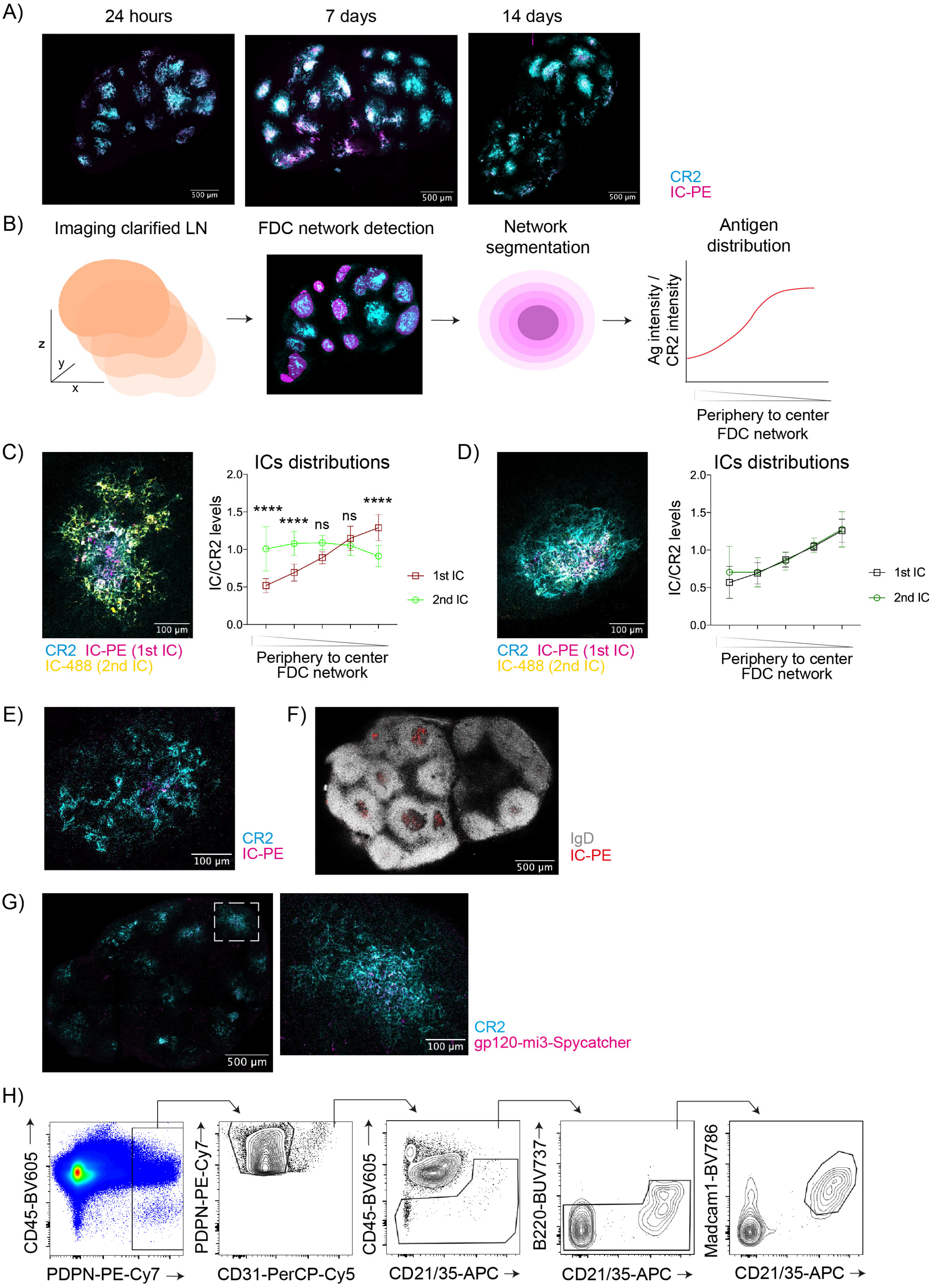

**Supplementary Figure 1. Antigen distribution on the FDC network centralizes over time and overlaps with GCs.**

A) Maximum intensity projection images of clarified LNs of mice immunized with IC-PE (magenta) at the indicated time points. FDC networks are shown in cyan (anti-CD21/35).
B) Image analysis workflow to quantify antigen distribution within each B cell follicle of clarified LNs. LNs stained with anti-CD21/35 antibodies to identify FDC networks are imaged in 3D by confocal microscopy. Volumes corresponding to the FDC networks of each follicle are segmented and then divided into concentric 3D shells. Antigen distribution is quantified as the fluorescence intensity of the antigen normalized by the fluorescence intensity of anti-CD21/35 staining in each shell and plotted according to the position of the shell from the periphery to the center.
C) A confocal image of an FDC network (cyan, anti-CD21/35) from a mouse immunized with two subsequent ICs in PBS analyzed 7 days after the first immunization ($1^{st}$ IC, IC-PE; magenta) and 24 hours after the second ($2^{nd}$ IC, IC-488; yellow). Right, graph shows the quantification of the distribution of both antigens (IC-PE, $1^{st}$ IC; IC-488, $2^{nd}$ IC) on the FDC network. (n = 8 LNs from 2 experiments).
D) A confocal image of an FDC network (cyan, anti-CD21/35) from a mouse immunized with two subsequent ICs in PBS analyzed 14 days after the first immunization ($1^{st}$ IC, IC-PE; magenta) and 7 days after the second ($2^{nd}$ IC, IC-488; yellow). Left graph shows the quantification of the distribution of both antigens (IC-PE, $1^{st}$ IC; IC-488, $2^{nd}$ IC) in the FDC network. (n = 12 LNs from 2 experiments).
E) A microscopy image of an FDC network (cyan, anti-CD21/35) from a LN of a mouse immunized with IC-PE (magenta) 56 days prior to imaging.
F) Confocal image of a draining LN 21 days after immunization with IC-PE (red). Naïve B cells are shown in grey (anti-IgD). IgD-negative areas within IgD-positive follicles indicate GCs.
G) A confocal image of a draining LN 7 days after immunization with AF555-labelled mi3-Spycatcher nanoparticles coated with YU-gp120-Spytag HIV envelope protein (magenta). FDC networks are shown in cyan (anti-CD21/35). White square indicates the region magnified.
H) Flow cytometry gating strategy to analyze FDCs.
Quantitative data shows the mean ± SD. Two-way's ANOVA with multiple comparisons, ns, $P > 0.05$; *, $P < 0.05$; **, $P < 0.01$; ***, $P < 0.001$

Supplementary Figure 2

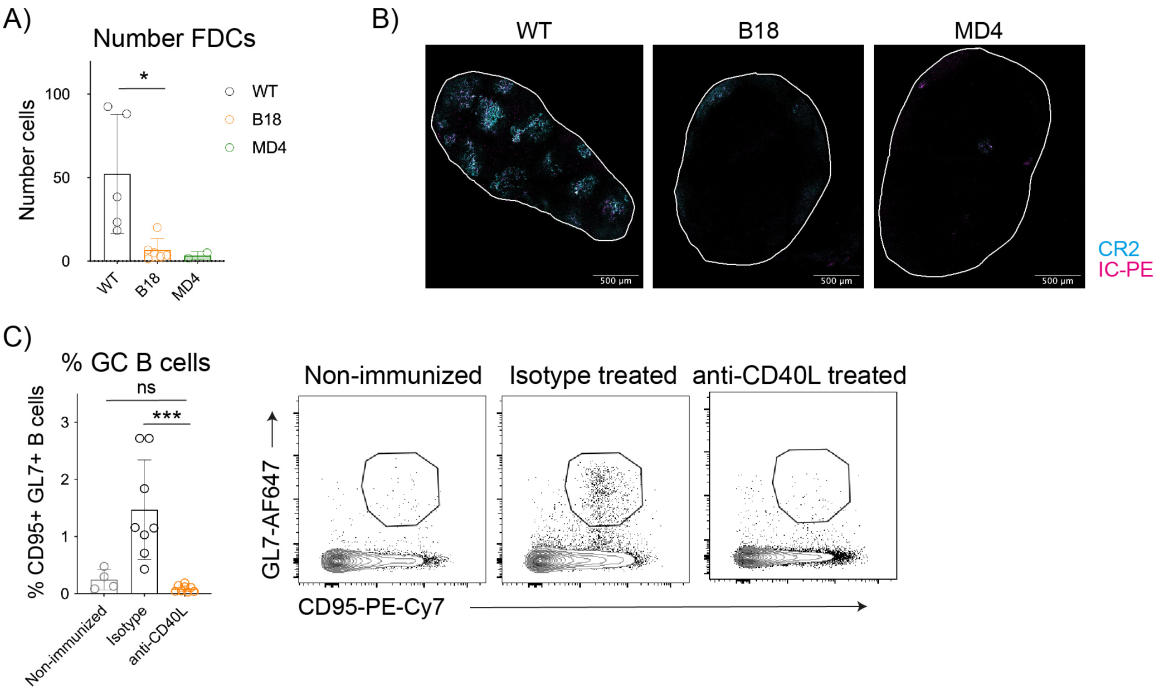


**Supplementary Figure 2. Antigen-specific B cell activation is required for FDC expansion.**

A) FDC numbers in non-transgenic C57BL/6 (WT) and BCR-transgenic B1-8f (B1-8flox IgK-/-) and MD4 mice 24 hours after immunization with IC-PE (n = 2-4 mice).

B) Representative confocal images of LNs from non-tg (WT), B1-8f and MD4 mice 24 hours after immunization with IC-PE (magenta). FDC networks are shown in cyan (anti-CD21/35). The white line delimits the edges of the organs.

C) Percentage of GC B cells in non-immunized (light grey) and IC-immunized mice treated with anti-CD40L (orange) or isotype control antibody (black) as described in Figure 2B (n = 4-8 mice, 2 experiments).
Quantitative data shows the mean ± SD. One-way ANOVA with multiple comparisons, ns, $P > 0.05$; *, $P < 0.05$; **, $P < 0.01$; ***, $P < 0.001$

Supplementary Figure 3

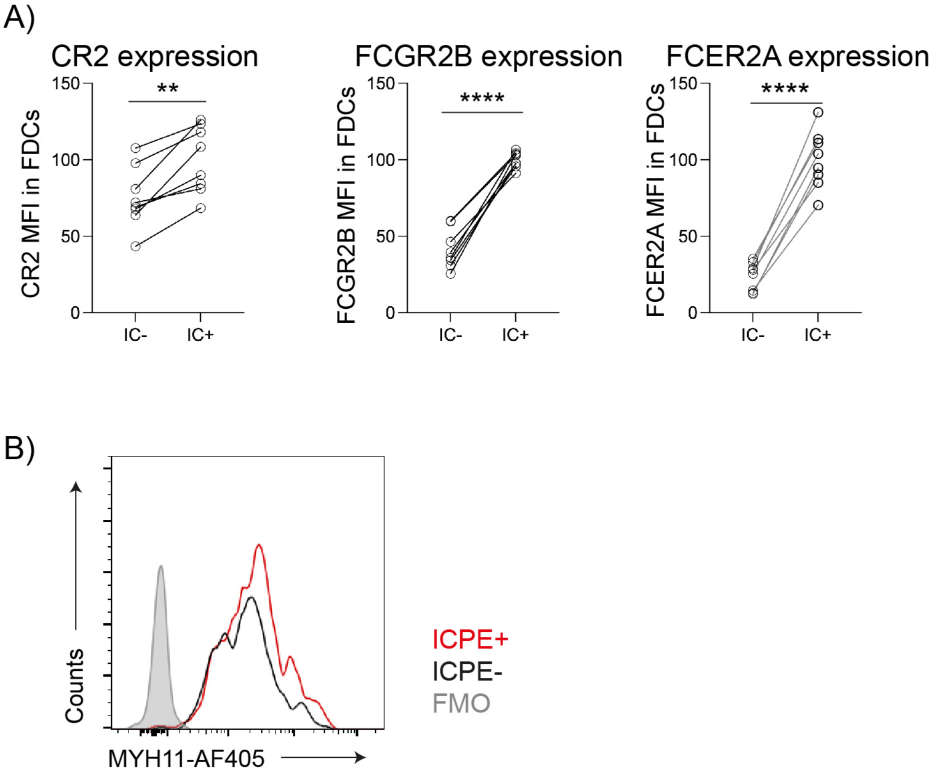


**Supplementary Figure 3. Central and peripheral LZ FDCs express different levels of IC-binding receptors on the membrane.**

A) CR2, FCGR2B and FCER2A membrane expression on $IC^{+}$ and $IC^{-}$ FDCs 7 days after immunization (n = 7 mice, 2 experiments).
B) Histogram showing Myosin heavy chain 11 (MYH11) expression in $IC\text{-}PE^{+}$ and $IC\text{-}PE^{-}$ FDCs 7 days after immunization.
Quantitative data shows the mean ± SD. Paired t-test, **, P < 0.01; ****, P < 0.0001

Supplementary Figure 4

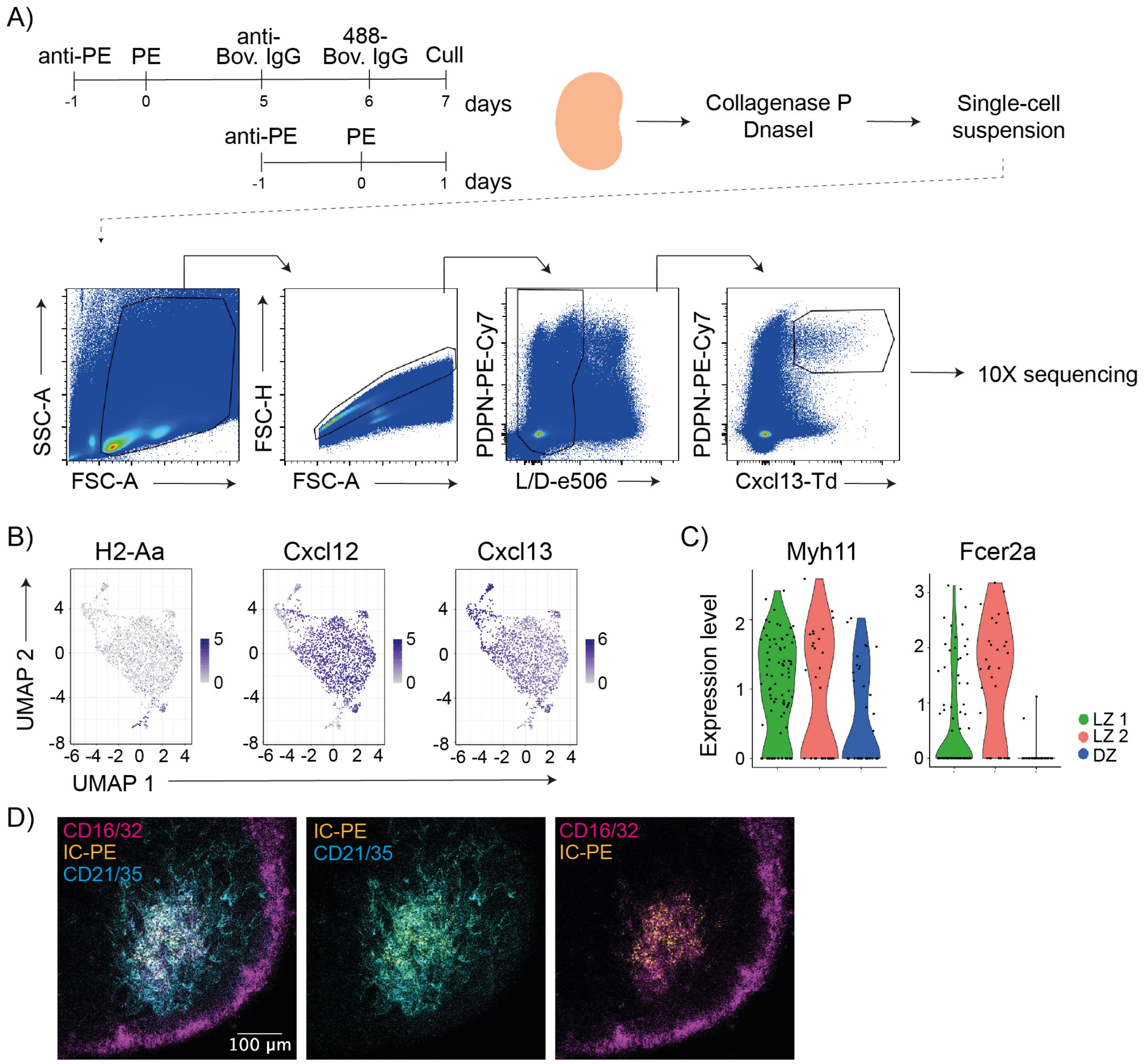


**Supplementary Figure 4. scRNAseq of follicular stromal cells.**

A) Experimental workflow for scRNAseq of Cxcl13-TdTomato$^{+}$ LN cells. Cxcl13-TdTomato mice were immunized consecutively with two ICs separated by 7 days only with one IC. 24 h after the last immunization, draining LNs were digested and dissociated in to a single-cell suspension that was stained. Live cells were flow-sorted based PDPN and TdTomato. Single sorted cells were used for 10x RNA sequencing.
B) Feature plots showing expression of markers for hematopoietic cells (H2-Aa) and cytokines important for LN organisation, Cxcl12 and Cxcl13.
C) Violin plots showing the expression of Myh11 and Fcer2a on the three FDC clusters from Figure 4C (LZ 1 in green, LZ 2 in red and DZ in blue).
D) Confocal image of a LN from a mouse after 7 days postimmunization with IC-PE (yellow). CR2 staining is shown in cyan and CD16/32 in magenta.

Supplementary Figure 5

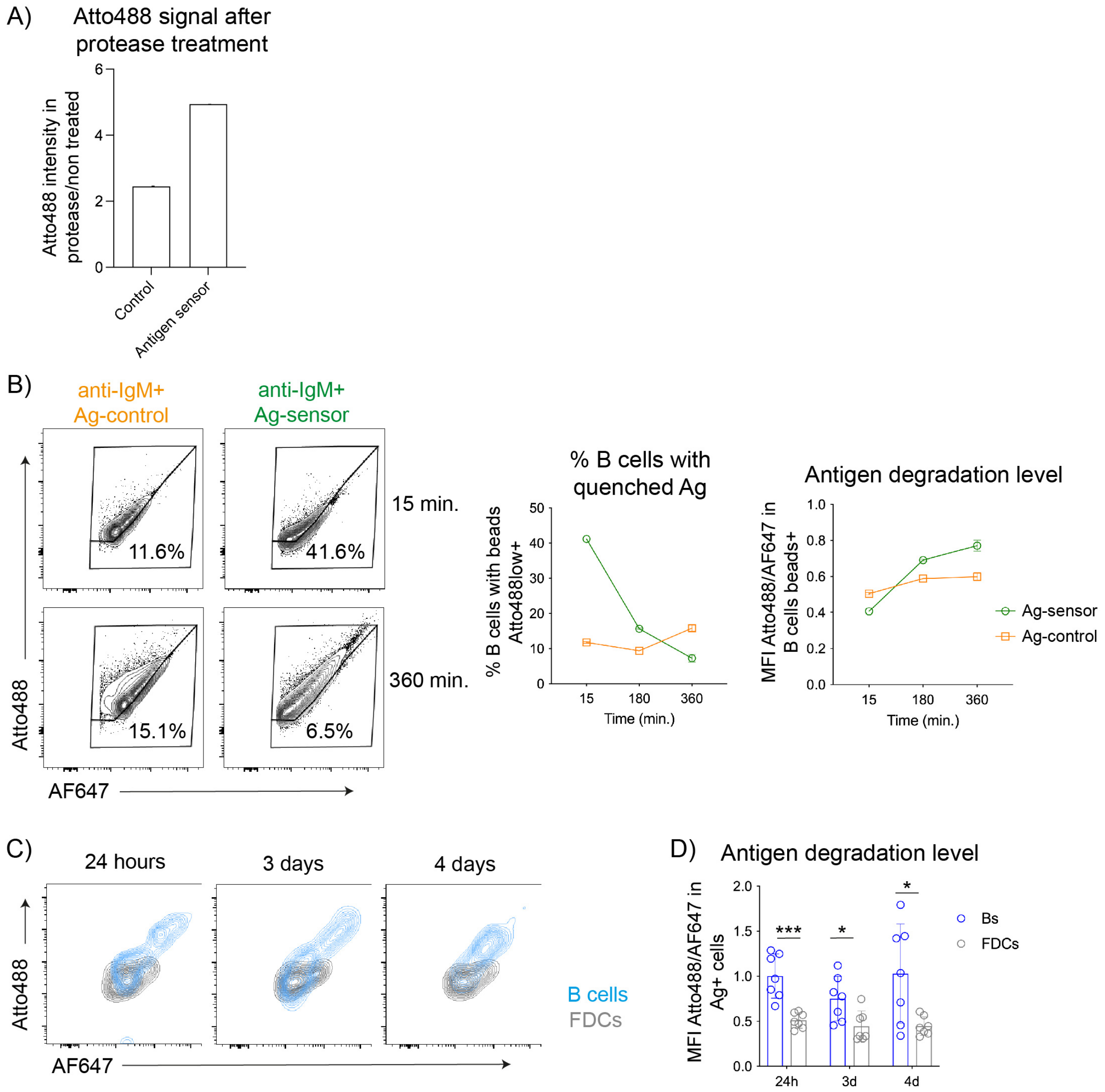


**Supplementary Figure 5. Antigen-sensor detects physiological antigen degradation.**

A) Quantification of Atto488 intensity after treating control sensor lacking the BHQ-1 quencher or antigen-sensor (1:9 antigen:quencher molar ratio) with protease for 30 min at 37C.
B) Naïve B cells were incubated with beads coated with anti-IgM and the antigen-sensor (green) or control sensor (orange) at indicated times at 37 °C. Plots illustrate Atto488 and AF647 intensity on B cells containing antigen-sensor beads or control sensor beads. Graphs show the percentage of B cells containing quenched antigen (%Atto488 low) and the levels of antigen degradation on B cells containing beads (measured as Atto488/AF647 intensity ratio).
C) Contour plots show Atto488 and AF647 levels on B cells (blue) and FDCs (grey) containing IC-antigen-sensor at different time points post-immunization.
D) Quantification of the antigen degradation levels in antigen$^{+}$ B cells and FDCs at different time points post immunization. (n = 7-8 mice, 2 experiments).
Quantitative data shows the mean ± SD. Unpaired t-test, ns, P > 0.05; *, P < 0.05; **, P < 0.01

Supplementary Figure 6

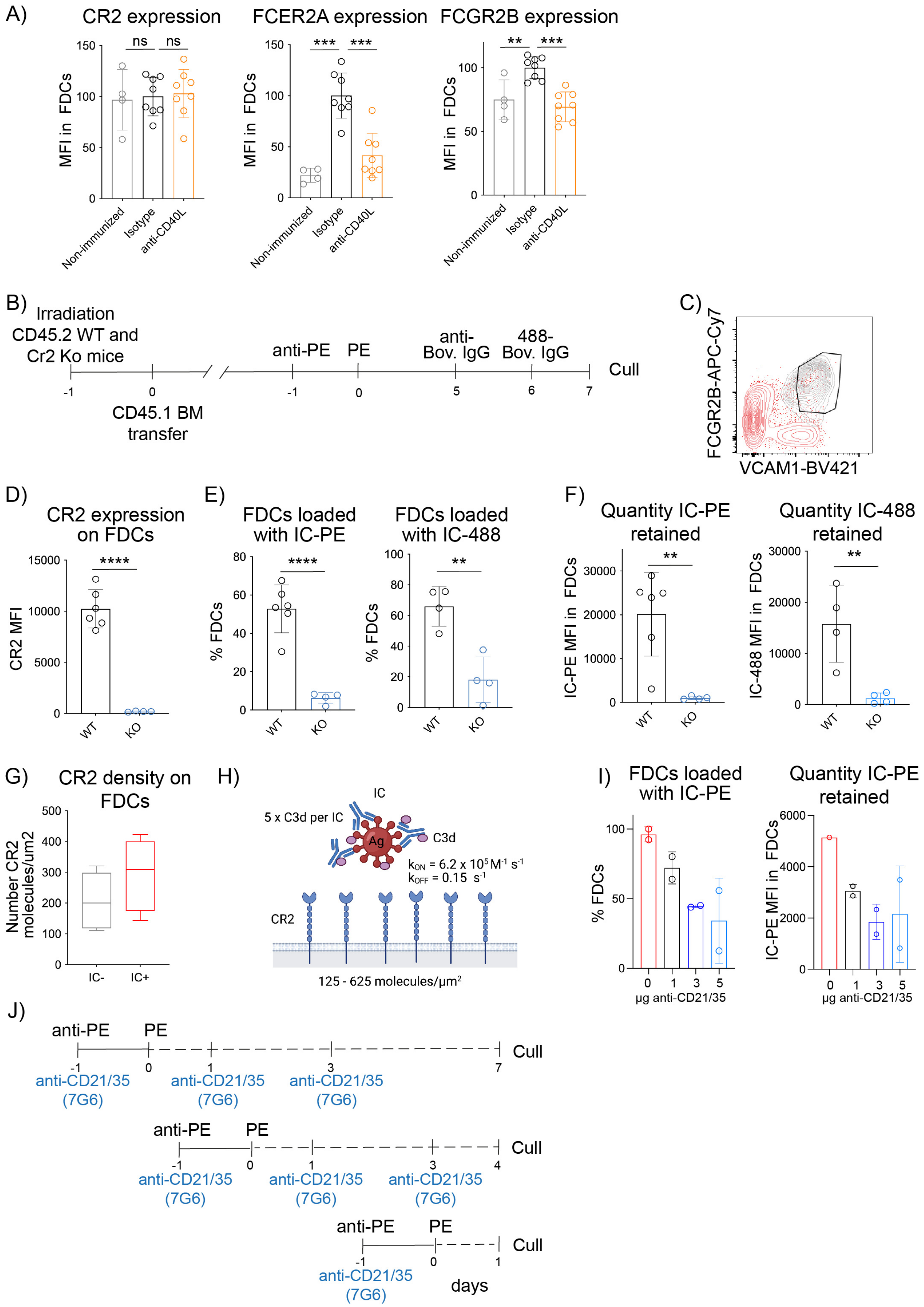

**Supplementary Figure 6. *In vivo* antigen IC deposition on FDCs requires CR2 expression.**

A) CR2, FCER2A and FCGR2B membrane expression on FDCs from non-immunized (grey) or immunized mice treated with anti-CD40L blocking antibody (orange) or isotype control antibody (black) as described in Figure 2C (n = 4-8 mice).
B) Experimental workflow. Lethally irradiated CD45.2 $Cr2^{+/+}$ (WT) and $Cr2^{-/-}$ (KO) mice were reconstituted with bone marrow cells from WT CD45.1/CD45.2 mice. 6 weeks after reconstitution, mice were immunized with IC-PE and again 6 days later with IC-488. LN were analyzed 24h after the last immunization.
C) Gating strategy to analyze FDCs in CD45.2 $Cr2^{+/+}$ (WT) and $Cr2^{-/-}$ (KO) mice reconstituted with WT CD45.1/CD45.2 BM. FDCs were identified based on the expression of FCGR2B and VCAM1 instead of CD21/35. In grey, FCGR2B and VCAM1 expression on WT FDCs ($PDPN^+$ $CD31^-$ $Madcam1^+$ $CD21/35^{hi}$). In red, FCGR2B and VCAM1 expression on $PDPN^+$ CD31- stromal cells.
D) CR2 expression on WT and Cr2 KO FDCs from BM reconstituted mice as described in (B) (n = 4-6 mice).
E) Percentage of FDCs loaded with IC-PE and IC-488 from mice described in (B) (n = 4-6 mice).
A) Quantity of IC loaded in the FDC network from mice described in (B) (n= 4-6 mice).
B) Quantification of surface CR2 density on $IC^+$ (red) or $IC^-$ (grey) FDCs 7 days after immunization. Quantibride PE beads were used as the molecular density standard (n = 8 mice, 2 experiments).
C) Schematic representation of the binding of a C3d-coated IC to the surface of an FDC illustrating the parameters for the mathematical modelling.
D) Quantification of the percentage of FDCs loaded with IC-PE and the amount (MFI) of IC-PE they were loaded with 24h after immunizing mice with IC-PE and injecting the indicated doses of anti-CD21/35 blocking antibody.
E) Immunization workflow used to analyse the effect of blocking CR2-C3d binding in vivo using an anti-CD21/35 antibody (7G6).

Quantitative data shows the mean ± SD. Unpaired t-test and One-way ANOVA with multiple comparisons, ns, P > 0.05; **, P < 0.01; ***, P < 0.001; ****, P < 0.0001

Supplementary Figure 7

A)

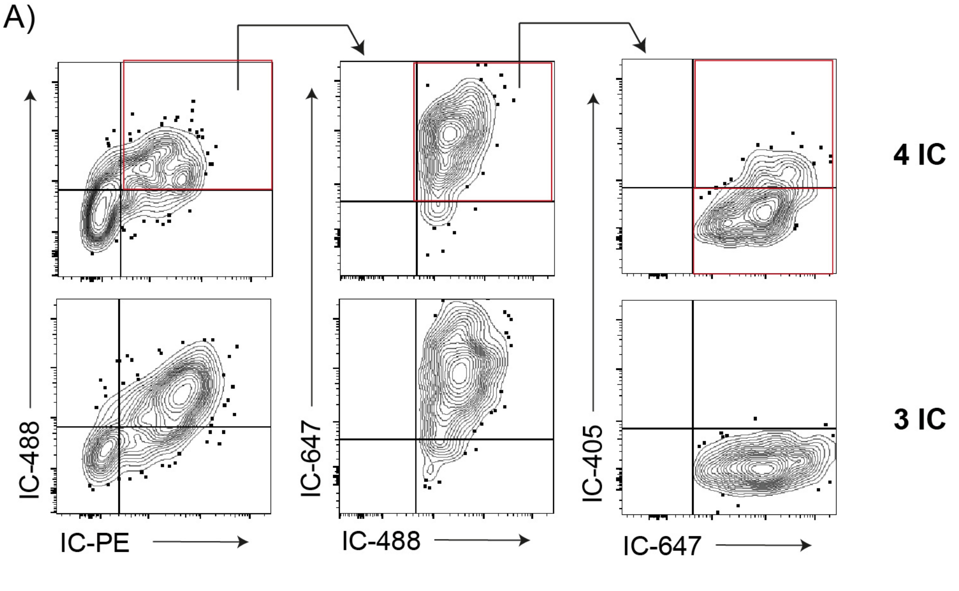


B)

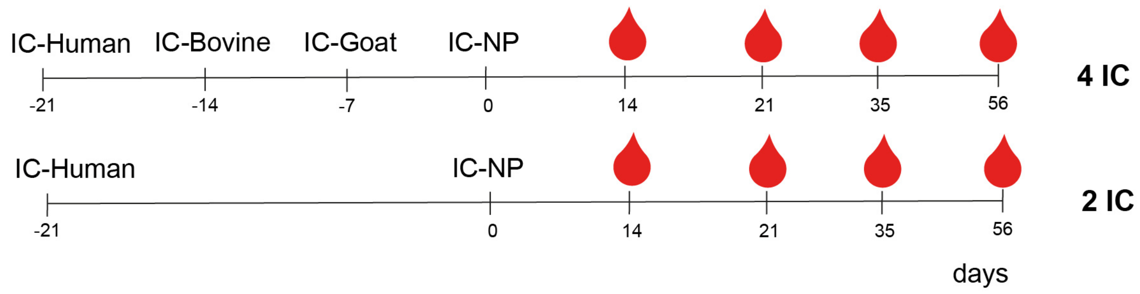


**Supplementary Figure 7. Central FDCs get partially saturated after consecutive antigen-IC immunizations, but still support and efficient antibody response.**

A) Flow cytometry gating strategy used to analyse FDCs containing different combinations of ICs from consecutive immunizations of mice immunized with 3 (lower panel) or 4 (upper panel) different fluorescent antigen-ICs as indicated in Figure 7A, B.

B) Immunization workflow followed to analyse the antigen-specific antibody response generated to NP under non-saturating (2-IC) or saturating conditions (4-IC).
